# Kinesin-4 Motor Teams Effectively Navigate Dendritic Microtubule Arrays Through Track Switching and Regulation of Microtubule Dynamics

**DOI:** 10.1101/2021.02.28.433181

**Authors:** Erin M. Masucci, Peter K. Relich, Melike Lakadamyali, E. Michael Ostap, Erika L. F. Holzbaur

**Affiliations:** Biochemistry and Molecular Biophysics Graduate Group, Perelman School of Medicine, University of Pennsylvania, Philadelphia, PA, 19104; Department of Physiology, Perelman School of Medicine, University of Pennsylvania, Philadelphia, PA, 19104

## Abstract

Microtubules establish the directionality of intracellular transport by kinesins and dynein through their polarized assembly, but it remains unclear how directed transport occurs along microtubules organized with mixed polarity. We investigated the ability of the plus-end directed kinesin-4 motor KIF21B to navigate mixed polarity microtubules in mammalian dendrites. Reconstitution assays with recombinant KIF21B and engineered microtubule bundles or extracted neuronal cytoskeletons indicate that nucleotide-independent microtubule binding regions of KIF21B modulate microtubule dynamics and promote directional switching on antiparallel microtubules. Optogenetic recruitment of KIF21B to organelles in live neurons resulted in unidirectional transport in axons but bi-directional transport with a net retrograde bias in dendrites; microtubule dynamics and the secondary microtubule binding regions are required for this net directional bias. We propose a model in which cargo-bound KIF21B motors coordinate nucleotide-sensitive and insensitive microtubule binding sites to achieve net retrograde movement along the dynamic mixed polarity microtubule arrays of dendrites.

## INTRODUCTION

Kinesins comprise a large superfamily of cytoskeletal motors that power transport in a polarized manner along microtubules (MTs) (Schnapp & Reese, 1989; Vale et al., 1985). Kinesin directionality is controlled by the organization of structurally polarized MT tracks (Erickson, 1974; Ludueña et al., 1977). Most kinesins move toward the more dynamic plus-ends of MTs, resulting in net outward, or anterograde, transport of cargos in cells with radially arrayed MTs. The mechanism of directional transport is more complicated in cell types in which the cytoskeleton is organized with mixed polarity (Muroyama & Lechler, 2017; Sanchez & Feldman, 2017). In mammalian neurons, MTs are organized with their plus-ends out within the soma and axon, but are oriented with mixed polarity in the dendrites (Baas et al., 1988; Heidemann & McIntosh, 1980). Intriguingly, the plus-end directed kinesin-4 KIF21B motor is associated with cargos moving predominantly in the retrograde direction in dendrites, despite the mixed polarity (Ghiretti et al., 2016). However, whether KIF21B is primarily responsible for retrograde directed motility and the mechanism by which KIF21B achieves this retrograde directionality is not known.

KIF21B is expressed in a variety of tissues (Marszalek et al., 1999) including the brain, where mutations are associated with developmental disorders (Asselin et al., 2020). In neurons, KIF21B localizes mainly to dendrites (Marszalek et al., 1999), although accumulating data suggest that KIF21B is important for cargo transport and the regulation of MT organization in both axons and dendrites (Asselin et al., 2020; Morikawa et al., 2018; Muhia et al., 2016). Mutations in KIF21B impede axon growth and branching, resulting in abnormalities in brain development and connectivity (Asselin et al., 2020), while the dendrites of neurons lacking KIF21B are less complex and exhibit tighter packing of MTs (Morikawa et al., 2018; Muhia et al., 2016). At the cellular level, KIF21B is involved in the endocytic recycling of NMDA receptors (Gromova et al., 2018), delivery of GABA_A_ γ2-subunits (Labonté et al., 2014) to the cell periphery, and the retrograde transport of TrkB-BDNF cargos (Ghiretti et al., 2016). Consistent with these roles in both cargo transport and MT organization, KIF21B-knockout mice display deficits in learning and memory (Muhia et al., 2016).

KIF21B interacts with MTs via a canonical N-terminal motor domain and secondary <u>MT</u> binding regions (MTRs) within the coiled-coil stalk and WD40 tail domains (Ghiretti et al., 2016; van Riel et al., 2017). Dimeric motor domain constructs lacking the C-terminal MTRs move processively toward the MT plus-end while the MTRs bind to MTs independently of the motor domain (Ghiretti et al., 2016; van Riel et al., 2017). The MTRs have been shown to dampen or pause MT dynamics (Ghiretti et al., 2016; Hooikaas et al., 2020; van Riel et al., 2017), but the magnitude of these effects differs depending on experimental conditions. A possible role for the MTRs in controlling directionality and processivity of cargo trafficking has not been explored.

In this study, we investigated how KIF21B motors establish long-range transport on mixed polarity MT networks. We started by examining KIF21B in simple *in vitro* systems, followed by *in vitro* reconstitution assays with added complexity, and finally live cell experiments using optogenetic motor recruitment to understand how KIF21B activity produces a net retrograde bias in neuronal dendrites. Consistent with a role in remodeling the MT cytoskeleton, recombinant KIF21B motors with intact MTRs stabilized MTs and promoted MT assembly at low nanomolar concentrations. The MTRs of KIF21B promoted track switching and long-distance transport on engineered antiparallel MT bundles *in vitro*; run length and directional switching were further enhanced by motor oligomerization. KIF21B motors were found to switch between anterograde and retrograde movement on stabilized native dendritic MT arrays prepared by cell extraction, but did not display the directional bias seen by KIF21B in live cells. These observations suggest that reconstitution of a directional bias requires MT remodeling by KIF21B motors, and thus would only be observed in motility along dynamic MT arrays. Using optogenetics to acutely recruit KIF21B to peroxisomes in live neurons, we found that KIF21B-bound cargos moved unidirectionally in axons but switched between anterograde and retrograde movement within dendrites; surprisingly, KIF21B recruitment induced net retrograde movement. We tested the requirements for native MT dynamics and the MTRs of KIF21B in live cell experiments, and found that both were required to achieve a net retrograde bias for the motility of dendritic cargos. Together these results suggest a mechanism where KIF21B teams bound to dendritic cargos coordinate both motor domains and MTRs to regulate MT dynamics and promote track switching to achieve long distance retrograde transport.

## RESULTS

### Recombinant KIF21B motors stabilize MTs and promote MT assembly

Live cell studies in neurons and T-cells indicate that KIF21B is involved in stabilizing MTs (Ghiretti et al., 2016; Hooikaas et al., 2020). However, *in vitro* experiments testing the effect of KIF21B on MT dynamics have produced different results depending on assay conditions (Ghiretti et al., 2016; van Riel et al., 2017). At low ionic strength and high motor concentrations, KIF21B increased MT dynamics by promoting faster growth speeds and catastrophe frequencies (Ghiretti et al., 2016). In contrast, experiments performed in higher ionic strength buffer in the presence of the end-binding protein EB3, resulted in the accumulation of KIF21B at the MT plus-ends and pronounced pausing of growth and shortening (van Riel et al., 2017). Given these differing results, our goal was to understand how KIF21B influences MT dynamics in a physiologically relevant system.

Here, we used Total Internal Reflection Fluorescence (TIRF) Microscopy and engineered system of single dynamic MTs to determine the ability of KIF21B motors to alter MT dynamics over a physiological range of motor concentrations (5 - 25 nM; Kim et al., 2014; Martens et al., 2006), in assays performed at physiological ionic strength. We compared the growth behavior of dynamic MT plus-ends in the absence or presence of Halo tagged full-length KIF21B (KIF21B-FL-Halo) or motor domain truncated KIF21B (KIF21B-MD-Halo) motors (Figure 1A-E and S1). The average MT growth rate decreased with increasing concentrations of either KIF21B-FL-Halo or KIF21B-MD-Halo (Figure 1F), although the magnitude of the effect was more pronounced in experiments with KIF21B-FL-Halo. Increasing concentrations of KIF21B-FL-Halo also induced a significant decrease in the catastrophe frequency, which was reduced to almost zero at KIF21B-FL-Halo concentrations above 5 nM. In contrast, addition of KIF21B-MD-Halo had no significant effect on the catastrophe frequency over all concentrations tested (Figure 1G). Of note, the motor domain of kinesin-1 (K560-Halo), known to stabilize MTs (Katsuki et al., 2014; Marceiller et al., 2005; Peet et al., 2018; Reuther et al., 2016) also suppressed the catastrophe frequency, but only at higher motor concentrations (≥ 25 nM); Figure S2). These results indicate that the KIF21B motor domain is sufficient to modulate MT dynamics, but inclusion of the C-terminal MTRs further enhances MT stabilization.

**Figure 1.**
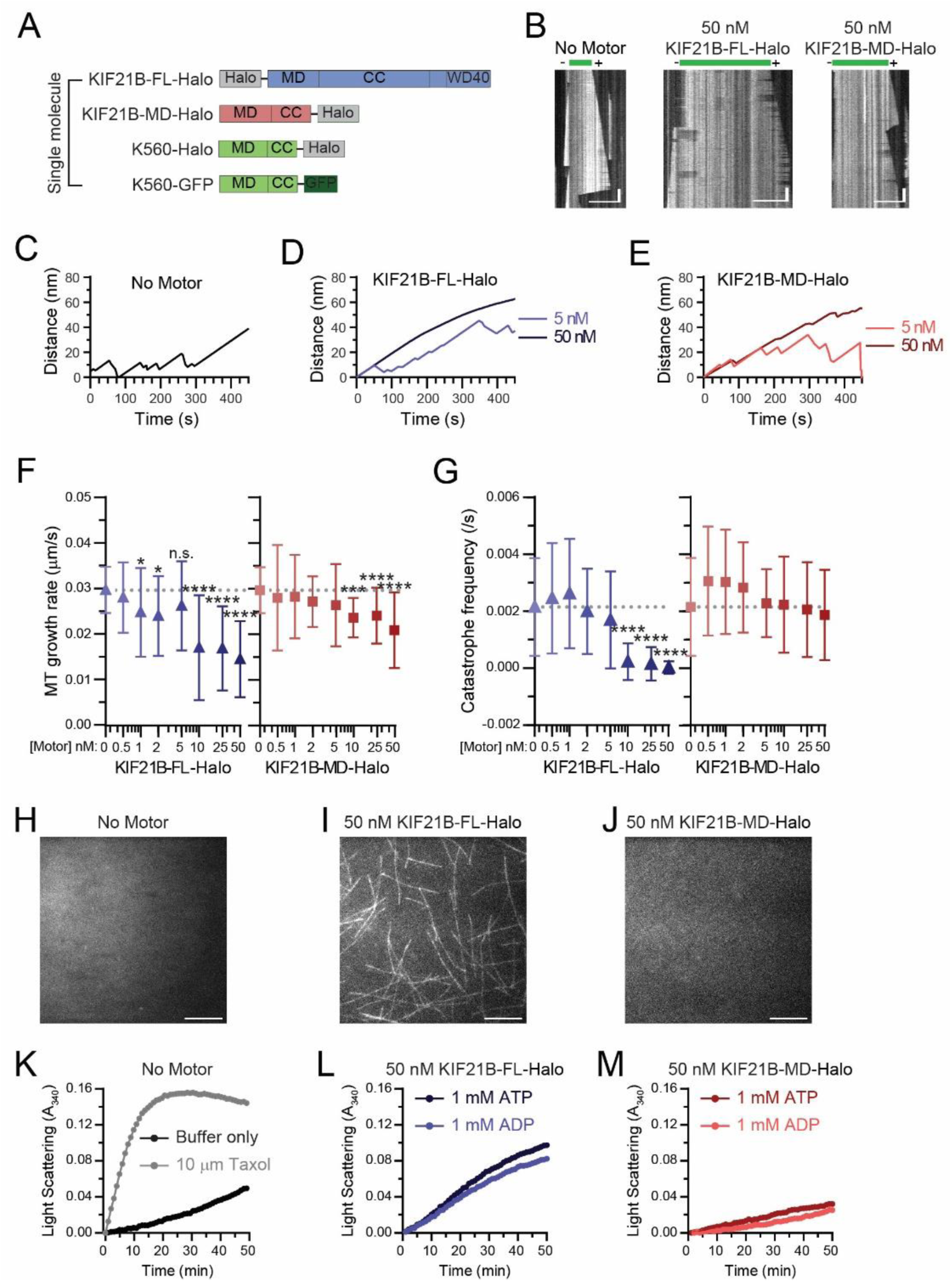
KIF21B motors stabilize MT dynamics and promote MT assembly. A) Motor constructs used for single molecule experiments. B) Kymograph plots showing dynamic MTs polymerizing from stabilized MT seeds in the absence or presence of KIF21B-FL-Halo or KIF21B-MD-Halo motors. Scale bars: 10 µm (horizontal) and 30 s (vertical). C-E) Average MT plus-end growth distances. Ten kymographs were averaged for each condition. N = 3 independent experiments. F) MT plus-end growth rates in the presence of increasing concentrations of KIF21B-FL-Halo or KIF21B-MD-Halo motors. Plotted are means and standard deviations. Kruskal–Wallis one-way ANOVA and Dunn’s multiple comparison (n.s. p > 0.05; *p < 0.05; ***p < 0.001; ****p < 0.0001). Data from 54-94 MTs and N = 3-6 independent experiments. G) MT plus-end catastrophe frequency in the presence of increasing concentrations of KIF21B-FL-Halo or KIF21B-MD-Halo motors. Plotted are means and standard deviations. Kruskal–Wallis one-way ANOVA and Dunn’s multiple comparison (****p < 0.0001). Data from 54-94 MTs and N = 3-6 independent experiments. H-J) TIRF microscopy images showing MT formation from free tubulin dimers incubated in the absence or presence of 50 nM KIF21B-FL-Halo or KIF21B-MD-Halo. Images were taken 10 min after solutions were introduced into flow chambers. Scale bar: 10 µm. K-M) Averaged light scattering traces for solutions containing buffer, Taxol, or 50 nM KIF21B-FL-Halo or KIF21B-MD-Halo motors with 1 mM ATP or 1 mM ADP. Means were compared with Kruskal–Wallis one-way ANOVA and Dunn’s multiple comparison (Buffer - Taxol ****p < 0.0001; Buffer – KIF21B-FL-Halo ****p < 0.0001; KIF21B-FL-Halo ATP – KIF21B-FL-Halo ADP n.s. p > 0.05; Buffer – KIF21B-MD-Halo n.s. p > 0.05; KIF21B-MD-Halo ATP – KIF21B-MD-Halo ADP n.s. p > 0.05). Data from 7-22 traces from N = 5 independent experiments. Graphs show mean values for each time point. See Figure S2 for graphs with standard deviations.

Differences with previous work (Ghiretti et al., 2016) likely result from improved motor preparations, lower motor concentrations, and use of higher ionic strength buffers that reduce the formation of KIF21B multimers and more closely model physiological conditions. We did not observe the prolonged attachment of KIF21B motors to dynamic MT plus-ends reported by van Riel et al. (2017), who performed experiments in the presence of mCherry-EB3. This suggests that both the reduction in MT growth rate and the decreased catastrophe frequency induced by full-length KIF21B may be mediated by a stabilization of the MT lattice, rather than a specific stabilization at dynamic MT plus-ends.

In our TIRF experiments, in addition to the dampened MT dynamics induced by KIF21B, we observed that new MTs emerged over time at 50 nM KIF21B-FL-Halo, which may be due to nucleation of new filaments. To determine if KIF21B motors promote the assembly of new MTs, we examined the effects of KIF21B on MT polymerization in the absence of preformed MT seeds. The addition of either buffer alone or KIF21B-MD-Halo was not sufficient to nucleate MTs, as assessed by TIRF microscopy and light scattering (Figure 1H-M and S3). However, addition of KIF21B-FL-Halo resulted in the robust formation of MTs over 10 min (Figure 1I), similar to the assembly induced by addition of K560-Halo (Figure S2F). The rate and extent of MT growth in the presence of KIF21B-FL-Halo, as assessed by turbidity, were similar in the presence of MgATP or MgADP (Figure 1L and S3G). Because the addition of MgADP weakens the interaction between kinesin motor domains and MTs (Crevel et al., 1996), but does not inhibit nucleation in our assays, these results suggest that the MTRs promote MT assembly. Collectively, the effects on MT growth rate, catastrophe frequency, and induction of assembly induced by full-length KIF21B lead to a net stabilization of MT growth at physiologically relevant concentrations of KIF21B, estimated to be 5 - 25 nM by mass spectrometry (Kim et al., 2014; Martens et al., 2006).

**Figure 3.**
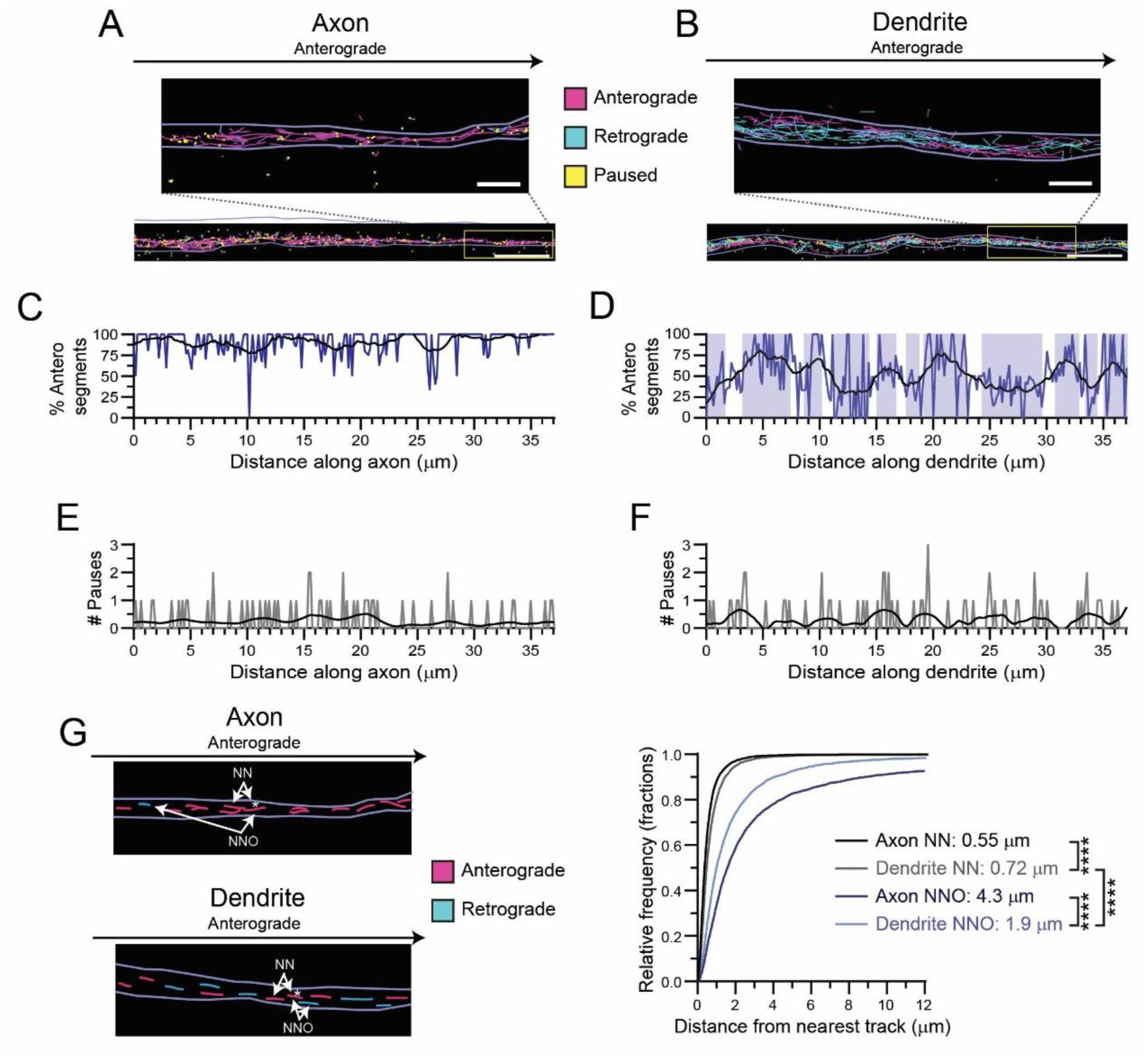
Dendritic MTs are distributed heterogeneously. A and B) Single particle motor track segments from a single TIRF movie. Light purple lines indicate bounds of axonal or dendritic process. Segments moving to the right (anterograde) are plotted in magenta, those moving to the left (retrograde) are cyan, and paused spots are shown in yellow. Zoom-in images show the separation between tracks. Scale bar: 5 µm. Zoom-in scale bar: 1 µm. C and D) Percentage of anterograde track segments across th e long axis of the axon and dendrite shown in A and B. Shaded blue areas indicate regions where the percentage of anterograde segments is above or below 50% in the dendritic process. Black lines show smoothed trendline over 2 µm window. E and F) Number of paused track segments located across the long axis of the axon and dendrite shown in A and B. Black lines show smoothed trendline over 2 µm window. G) Schematic of nearest neighbor (NN) track and nearest opposing neighbor (NNO) track, and cumulative distribution of the distance between each anterograde moving track segment and its NN moving in any direction or NNO moving in the opposite direction in both axons and dendrites. A single exponential decay function was fit to the data. Listed are the average distance traveled, as calculated by taking the inverse of the rate constant. Kruskal–Wallis one-way ANOVA and Dunn’s multiple comparison (****p < 0.0001). Unless otherwise indicated, data from 59-80 processes from n = 49-50 neurons and N = 4 independent experiments.

### Reconstitution of motor switching on engineered parallel and antiparallel MT bundles

Since KIF21B motors can effectively navigate the antiparallel MT arrays characteristic of dendrites in mammalian neurons (Ghiretti et al., 2016), we wanted to test if the relative orientation of MTs within MT bundles affects the motile properties of KIF21B. As KIF21B-FL motors modulate MT dynamics (Figure 1), we reconstituted parallel and antiparallel arrays of dynamic MTs (Figure 2A). The majority of the bundles were composed of two aligned MTs, oriented either parallel or antiparallel to one another, as evaluated by kymograph analysis of MT plus-end dynamics and fluorescence intensity measurements over time (Figure 2A and Video 1-3).

**Figure 2.**
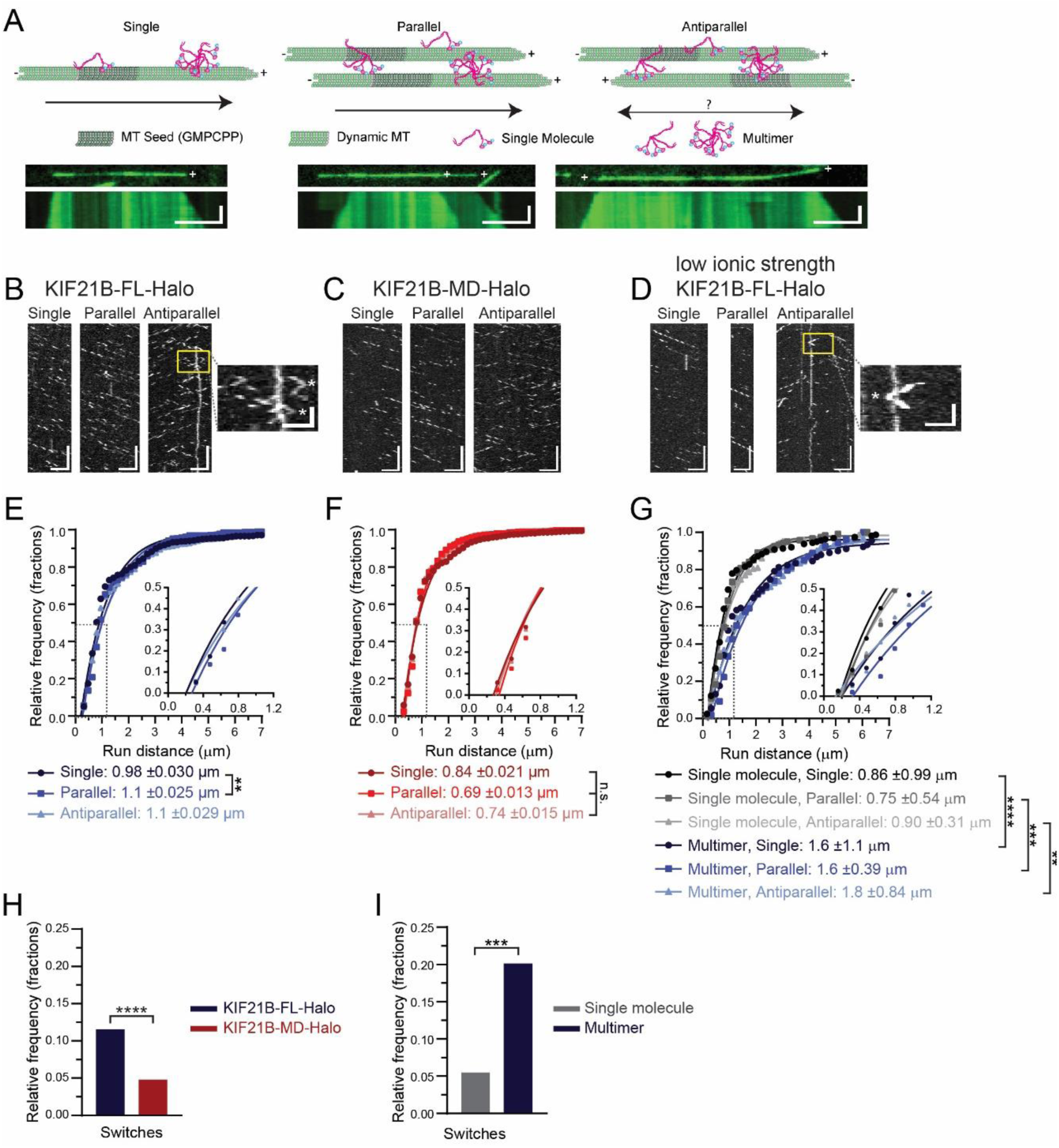
Motor multimerization and KIF21B MTRs improve motor processivity and track switching. A) Schematic of *in vitro* single molecule assays. Dynamic MTs were grown from surface-immobilized GMPCPP MT seeds. Some MTs remain isolated in space as they grow, while others overlap in parallel or antiparallel orientations with neighboring MTs. Kymographs show growing dynamic MT ends over time for each type of MT arrangement. Dynamic MT plus-ends are indicated (+). KIF21B dimers or multimeric complexes contact MT lattices with their N-terminal motor domains, as well as MTRs within their C-terminal tails. Motor domains were labeled with TMR Halo ligands (cyan dot). Scale bars: 5 µm (horizontal) and 1 min (vertical). B-D) Kymograph plots of KIF21B-FL-Halo, KIF21B-MD-Halo and KIF21B-FL-Halo motility in low ionic strength buffering system along single MTs or parallel and antiparallel MT bundles. Zoom-in image highlights track switches observed on antiparallel MTs (indicated with white * in expanded images). Track switches were defined as a reversal in run direction exceeding 3 pixels. Scale bars: 5 µm (horizontal) and 20 s (vertical). Zoom-in scale bars: 2.5 µm (horizontal) and 5 s (vertical). E-G) Cumulative distribution of total distance motor traveled along different MT arrays. Data points were fitted by a single exponential decay function. Differences in run distances can be more clearly seen in insets. Average distance traveled, as calculated by taking the inverse of the rate constant, and standard deviations are shown in the legends. Kruskal–Wallis one-way ANOVA and Dunn’s multiple comparison (n.s. p > 0.05; **p < 0.01; ***p < 0.001; ****p < 0.0001). H and I) Frequency of track switches for single KIF21B-FL-Halo and -MD-Halo motor runs or single and multimeric KIF21B-FL-Halo motor run along antiparallel MTs. Two-tailed Mann-Whitney (***p < 0.001; ****p < 0.0001). Data for single molecule KIF21B-FL-Halo and KIF21B-MD-Halo from 400-1069 motor runs from n = 20-60 MTs and N = 6-7 independent experiments. Data for single and multimeric KIF21B-FL-Halo from 54-174 motor runs from n = 11-12 MTs and N = 4 independent experiments.

We compared the motility of purified recombinant KIF21B-FL-Halo or -MD-Halo motors (Figure 1A) on single MTs and MT bundles organized in either parallel or antiparallel orientations, as assessed by monitoring the differing assembly and disassembly dynamics characteristic of MT plus- and minus-ends (Figure 2A-D, Video 1-3). We focused on analyzing only motor particles whose intensity was consistent with single molecules, of which ∼ 50% were fully labeled. Both the full-length and motor domain constructs moved uniformly in the plus-end direction on single MTs (Figure 2B). We observed robust unidirectional movement on parallel MT bundles (Figure 2B and C), and robust bi-directional movement on antiparallel MT bundles (Figure 2B and C). Both KIF21B-FL-Halo and -MD-Halo motors moved with similar average speeds along the three MT arrangements (Figure S4A and B). However, KIF21B-MD- Halo displayed shorter run distances than full-length KIF21B-FL-Halo, regardless of the MT arrangement (Figure 2E and F). The run distance of KIF21B-FL-Halo was slightly, but significantly, longer on parallel MT bundles compared to single MTs, while the run distance of KIF21B-MD-Halo was not affected by MT track type. Strikingly, both KIF21B-FL-Halo and -MD-Halo motors exhibited track switching within antiparallel bundles (Figure 2B zoom-in). Switching was more frequent for the full-length construct, measured as either total events observed (Figure 2H) or number of switches per distance traveled (Figure S4C).

In the cell, cargos are likely to be transported by more than one kinesin motor (Gross et al., 2007; Holzbaur & Goldman, 2010). Multimerization of KIF21B may also have been observed by van Riel et al., (2017), where KIF21B motor complexes formed at the plus-ends of growing MTs. To test the effects of motor multimerization on motility and track switching, we introduced purified KIF21B-FL-Halo motors into low ionic strength buffer (12 mM PIPES, pH 6.8), which induced multimer formation. We used single molecule photobleaching to calibrate the relationship between fluorescence intensity and the number of molecules in a cluster (Figure S5A) and used this calibration to distinguish between single molecules and multimers (estimated to be 2-3 motors) (Figure S5B). KIF21B-FL-Halo multimers and single molecules displayed similar velocities on all MT arrays (Figure S5D), but multimers moved longer distances than single molecules regardless of the type of MT arrangement (Figure 2G). Run distances positively correlated with intensity while velocity did not correlate with intensity, suggesting that formation of motor teams leads to improved run distances with no effect on velocity (Figure S6A and B).

KIF21B-FL-Halo multimers switched tracks significantly more frequently per run than single molecules of KIF21B-FL-Halo (Figure 2I); multimeric motors also switched tracks more frequently per unit distance than single molecules (Figure S5E). However, we did not observe a correlation between the number of track switches per distance traveled and the intensity of the oligomer (Figure SC), suggesting that multimerization of only two motors is needed to promote track switching. Collectively these results indicate that the MTRs in the C-terminal of KIF21B enhance both processivity and track switching, while motor multimerization further contributes to both of these parameters.

### Quantitative mapping of the dendritic MT network

We observed that bundles of only two MTs are sufficient to influence KIF21B motility. How does this scale to bundles of MTs organized into larger arrays, such as found in dendrites? Live cell imaging studies suggest that KIF21B drives endogenous cargos with a net retrograde bias in dendrites (Ghiretti et al., 2016), a puzzling finding for a plus-end directed motor moving along an array in which the majority (∼ 60%; Ayloo et al., 2017) of MTs are oriented with their plus-ends outward. A recent model proposed by Kapitein and colleagues (Tas et al., 2017) suggests that the dendritic MT network is composed of distinct unipolar MT bundles, and that these bundles are marked by specific post-translational modifications (PTMs) of the tubulin cytoskeleton. Tas et al., (2017) proposed that MT motors recognize this code, preferentially binding to and moving along selected MT populations to drive long-range transport in a specific direction. Does the MT code present in dendrites bias KIF21B motors to move in the retrograde direction?

To determine if the native MT arrays in dendrites can bias motors toward movement in the retrograde direction, we used motor-PAINT with rat hippocampal neurons (Brawley & Rock, 2009; Tas et al., 2017), which allows super-resolution tracking of single-motors interacting with the cytoskeleton of fixed permeabilized cells. We used immunocytochemistry to compare the ratios of tyrosinated and acetylated tubulin intensities in axons and dendrites on extracted arrays compared to permeabilized neurons and found no significant differences (Figure S7), indicating that these tubulin PTMs are preserved after extraction.

To extend the previous work of Tas et al., (2017) and generate a more quantitative map of the MT network within hippocampal dendrites, we used TIRF microscopy to track individual processive runs of K560-GFP (Figure 1A) along native MT arrays extracted from axons and dendrites (Figure 3A and B, Video 4 and 5), with the assumption that this motor provides a non-biased readout of MT orientation. The resulting videos demonstrate the expected pattern of primarily unidirectional motility along axons and more bi-directional motility along dendrites, but the density of the native MT arrays makes unambiguous run detection more challenging. To more rigorously map the runs, we used an automated single particle segmentation and tracking program called Cega that we developed to track moving particles within a noisy environment (Masucci, Relich et al., 2020; see Methods). Due to the crossing of individual motor runs in both axons and dendrites, complete tracks could not be reliably determined. Instead, individual track segments from single motors connected frame-by-frame from motor movement captured every 0.2 s for 2 min were identified and used to construct a quantitative map of MT organization and polarity.

Cega analysis demonstrates that K560-GFP motors moved along axonal MT arrays with > 90% of the tracks in the anterograde direction (Figure 3A and C, Video 4). On dendritic MT arrays, K560-GFP motors moved bi-directionally (Figure 3B and D, Video 5), but strikingly, some regions showed preferred directionality as indicated by blue shading in Figure 3D; this local heterogeneity in MT organization is also apparent from the smoothed trend line that shows the average track direction within 2 µm (compare the trend lines from axons and dendrites in Figure 3C and D). In contrast to the localized differences in MT orientation, pauses were randomly distributed throughout both axons and dendrites (Figure 3E and F and S8).

To investigate the spatial relationship of adjacent MT tracks, we measured the distance between anterograde moving track segments and their nearest neighboring (NN) track segment or nearest neighboring opposing (NNO) retrograde track segment in both axons and dendrites (Figure 3G). The distance between NN track segments in axons was significantly shorter than that of dendrites, consistent with tighter MT spacing in axons (J. Chen et al., 1992). If the dendritic MTs were homogenously mixed in orientation, then the distance between the anterograde track segments and their NN and NNO track segment should be similar. However, in both axons and dendrites, the distance between NN track segments was much shorter than that of NNO track segments. In axons, this is likely due to the very small number of MTs oriented with plus ends towards the soma (Baas et al., 1988). In dendrites, the increased NNO as compared to NN distance provides quantitative support for the model proposed by Tas et al. (2017) which suggests that MTs are locally organized in bundles with similar polarities, creating regions where similar oriented MTs are located close together and those of opposite orientation are spatially removed. Further, our observations indicate that MT orientation in mammalian dendrites is distributed heterogeneously along the lengths of each process.

### Recombinant KIF21B motors exhibit directional switches on extracted dendritic MT arrays

To determine if the native MT code together with the native organization of MTs in dendrites leads to a retrograde bias for KIF21B motility, we used the motor-PAINT assay to observe the movement of KIF21B molecules in axons and dendrites (Figure 4 and S9). We were able to directly compare the movement of either Alexa Fluor 660-labeled KIF21B-FL-Halo or KIF21B-MD-Halo to K560-GFP (Figure 1A) by mixing the differentially labeled motors and simultaneously tracking their movement on the same MT arrays (Figure 4A and B). Individual runs along extracted MT arrays were tracked using Cega (Masucci, Relich et al., 2020). Kymographs generated for the KIF21B constructs show that both the full-length and truncated motors have similar directional patterns along axonal and dendritic MT arrays, and both closely resembled the patterns seen with K560-GFP (Figure 4A and B). Again, we saw evidence for regional heterogeneity in the organization of MTs in dendrites, with some kymographs demonstrating motors moving in mainly one direction (Figure 4A “Bi-directional motility” versus “Anterograde motility”), supporting the idea that the MTs are locally organized into bundles with similarly oriented MTs (Tas et al., 2017).

**Figure 4.**
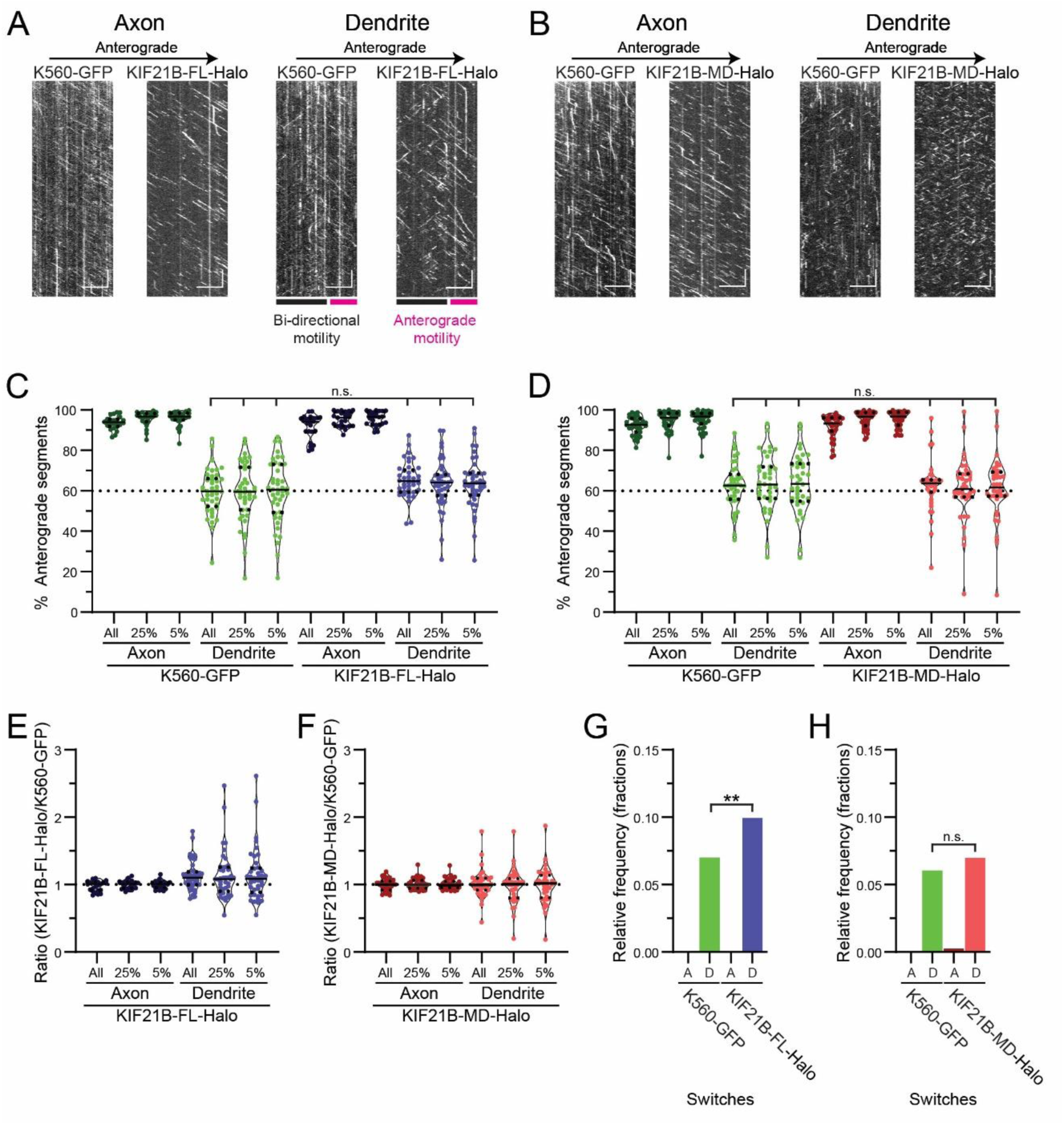
KIF21B motors do not move with a directional bias on extracted dendritic MT arrays. A and B) Kymographs of motor motility along axonal or dendritic MTs for K560-GFP and KIF21B-FL-Halo or KIF21B-MD-Halo motors moving on the same MTs. Colored bars in Figure 4A indicate regions of the dendrite where motors move bi-directionally, and regions where motors move primarily in one direction (anterograde motility). Scale bars: 10 µm (horizontal) and 10 s (vertical). C and D) Percentage of track segments moving in the anterograde direction in axons and dendrites. Plotted are individual values in color and means in solid black lines and 25^th^ and 75^th^ quartiles in dotted black lines. Kruskal–Wallis one-way ANOVA and Dunn’s multiple comparison (n.s. p > 0.05). E and F) Ratio of the percentage of KIF21B-FL-Halo or KIF21B-MD-Halo anterograde track segments to the percentage of K560-GFP anterograde tracks along the same axonal and dendritic arrays. Plotted are individual values in color and means in solid black lines and 25^th^ and 75^th^ quartiles in dotted black lines. G and H) Frequency of motors switching direction of movement along axonal (A) and dendritic (D) MT arrays. Kruskal–Wallis one-way ANOVA and Dunn’s multiple comparison (n.s. p > 0.05; **p < 0.01). Data from 24-42 processes from n = 21-28 neurons and N = 4 independent experiments.

Along axonal MT arrays, > 90% of all motors tested moved in anterograde direction, while ∼ 60% moved in the anterograde direction on dendritic MTs (Figure 4C and D). The standard deviations of directional segments in dendrites for all three motors were much larger than observed in axons, again consistent with local variability in MT organization along the length of the dendrite. We found no difference in direction of movement between K560-GFP and KIF21B-FL-Halo or KIF21B-MD-Halo in axons and dendrites (Figure 4C-F). We further compared the percent anterograde direction of KIF21B- FL-Halo and KIF21B-MD-Halo motors in dendrites to 60%, since this is the direction of motility expected from the 60% plus-end out MT polarity measured by K560-GFP motors, and measured no significant difference. This comparison suggests that native, stabilized dendritic MT arrays do not bias KIF21B motors to move in the retrograde direction.

Our preparations of purified K560-GFP, KIF21B-FL-Halo, and KIF21B-MD-Halo motors produced mainly single, two-headed motor proteins, as assessed by controlled photobleaching of processing motors in motility assays (Figure S5A and B). However, ∼ 40% of the population of KIF21B-FL-Halo motors were multimers, as determined by fluorescence intensity measurements. We took advantage of this multimerization to test the idea that multiple motors are needed to confer retrograde bias to KIF21B. We compared the percentage of anterograde segments for each process for all motors to that of the brightest 25% or 5% of moving motors and found no significant difference between these groups and no significant difference against the 60% expected motility direction (Figure 4C and D). We further quantified the ratio of the percentage of anterograde track segments of KIF21B motors compared to K560-GFP motors for each process, and found that the ratio of KIF21B to K560-GFP segments was about 1 (Figure 4E and F), regardless of the motor brightness, suggesting that motor multimerization does not contribute to directional bias in motility along extracted neuronal MT cytoskeletons.

In dendrites but not axons, we observed pronounced directional switching by KIF21B-FL-Halo. We measured the frequency of motor runs that switched direction of movement along dendritic and axonal MT arrays, and observed that KIF21B-FL-Halo switched tracks more frequently than K560-GFP along the same dendritic MT tracks (Figure 4G). In contrast, KIF21B-MD-Halo and K560-GFP switched tracks at a similar rate (Figure 4H). These results suggest that the MTRs in the C-terminal tail of KIF21B enhance track switching along dendritic MT bundles with native organization and spacing.

### Optogenetic recruitment of KIF21B induces retrograde trafficking of dendritic cargos in live neurons

As the native MT code and cytoskeletal organization of extracted dendrites was insufficient to establish a retrograde bias for KIF21B motors that paralleled that observed in live cell assays (Ghiretti et al., 2016), we wondered whether the key factor missing from our assay might be MT dynamics. Previous work (Ghiretti et al., 2016; Hooikaas et al., 2020; van Riel et al., 2017) and our current studies indicate that KIF21B has the ability to modulate MT dynamics both *in vitro* and *in vivo*. This modulation may contribute to the establishment of a net retrograde bias for KIF21B-driven cargos moving along mammalian dendrites as KIF21B could influence the dynamics of specific populations of MTs oriented plus-end in over those oriented plus-end out based on differences in stability. Because of this, active MT remodeling by KIF21B could be required to generate retrograde biased motility observed in live cell assays (Ghiretti et al., 2016).

To test this possibility, we used a previously validated optogenetic assay (Ayloo et al., 2017) to determine how KIF21B motors perform when recruited to a non-native cargo in axons and dendrites of live neurons, under cellular conditions with robust MT dynamics (Figure 5A). Full-length KIF21B (KIF21B- FL) was recruited to peroxisomes, a largely immobile cargo in hippocampal neurons, via the heterodimerization of DHFR-fused motor and HaloTag (Halo)-fused peroxisome targeting sequence PEX3 in response to the photo-induced uncaging of a small organic compound (CTH) (Ayloo et al., 2017; Ballister et al., 2014, 2015; Zhang et al., 2017). The CTH was uncaged in defined regions of the dendrites and axons of neurons expressing both peroxisome and KIF21B constructs. Photoactivation induced the recruitment of mCherry-labeled KIF21B to peroxisomes labeled with GFP (Figure 5B-C, Video 6 and 7) and induced the processive motility of more than 60% of peroxisomes in axons and dendrites within ∼ 28 s (Figure 5D, S10).

**Figure 5.**
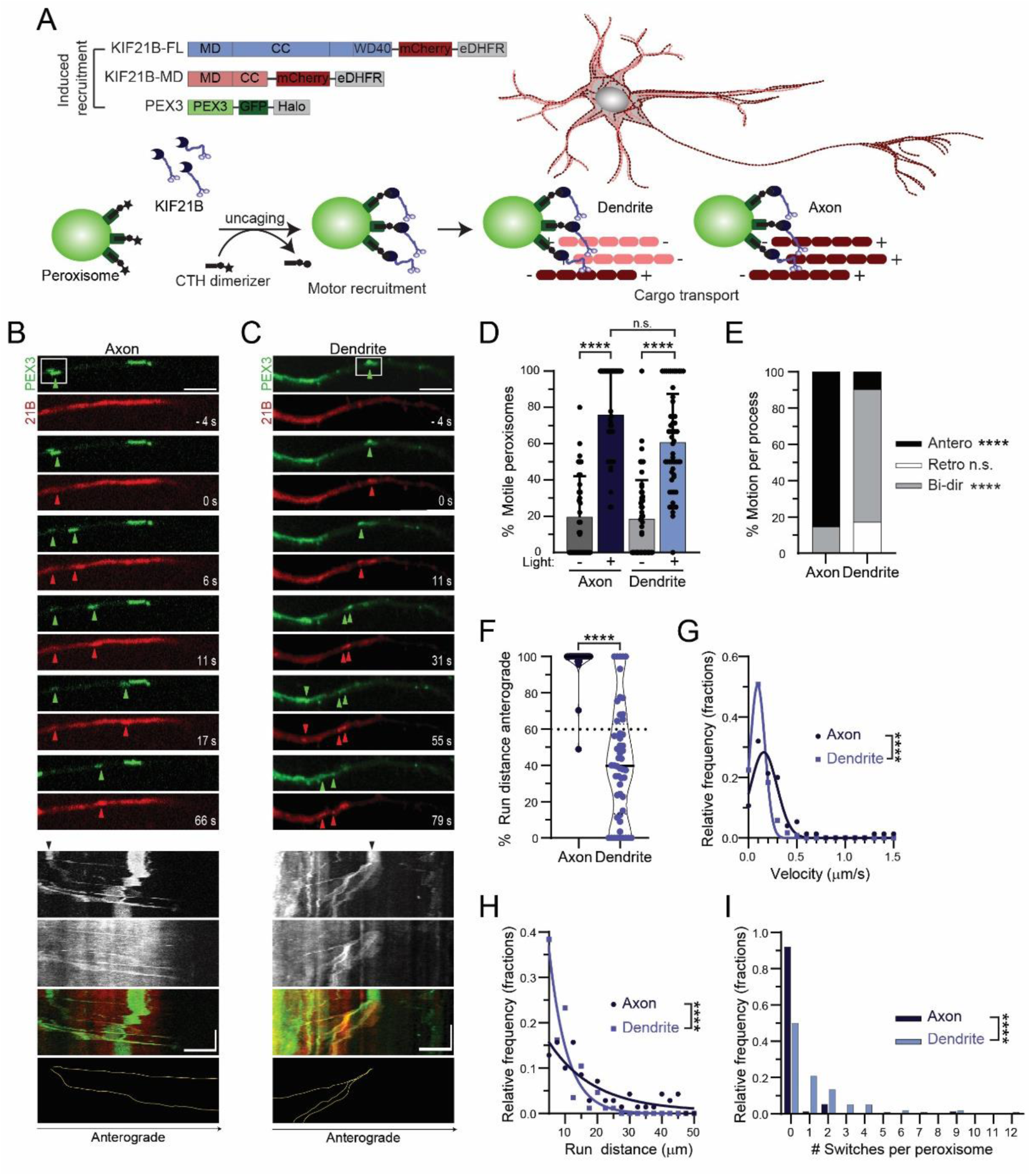
Induced recruitment of KIF21B motors to peroxisome cargo in live neurons promotes retrograde trafficking in dendrites. A) Diagram of motor constructs and experimental schematic for induced recruitment experiments in live neurons. Dark red and pink lines and plus and minus signs indicate MT orientation within arrays; antiparallel MT arrays in dendrites and parallel arrays in axons. B and C) Time series and corresponding kymograph showing movement of photoactivated (white box) peroxisomes in axons and dendrites. Green and red arrowheads indicate the locations of photoactivated peroxisomes with successful KIF21B recruitment in both the PEX3 and KIF21B fluorescence channel, respectively. Black arrows on kymographs indicate regions of photoactivation. Yellow traces indicate movement of the KIF21B recruited peroxisomes identified in the time series. Scale bars: 5µm (horizontal) and 1 min (vertical). D) Percentage of peroxisomes that are motile in axons and dendrites with and without photoactivation. Plotted are means and standard deviations. Kruskal–Wallis one-way ANOVA and Dunn’s multiple comparison (n.s. p > 0.05; ****p < 0.0001). E) Percentage of peroxisome runs moving anterograde, retrograde or bi-directionally for each axonal and dendritic process. Bi-directional (Bi-dir) movement is characterized by peroxisomes that move more than 0.4 µm in both anterograde (Antero) and retrograde (Retro) directions. Two-way ANOVA and Tukey multiple comparison (n.s. p > 0.05; ****p < 0.0001). F) Percentage of total peroxisome run distance per process moving in the anterograde direction. Plotted are individual values in color, means in solid black line and 25^th^ and 75^th^ quartiles in dashed black lines. Two-tailed Mann-Whitney test (****p < 0.0001). G) Histogram of velocities of motile peroxisomes after photoactivation in axons and dendrites. Data points were fitted by a gaussian function. Two-tailed Mann-Whitney test (****p < 0.0001). H) Histogram of un distances of motile peroxisomes after photoactivation. Data points were fitted by a single exponential decay function. Two-tailed Mann-Whitney test (****p < 0.0001). I) Histogram of number of track switches per peroxisome run in axons and dendrites. Track switches are characterized by a reversal in direction that exceeds 0.4 µm in both anterograde and retrograde directions. Two-tailed Mann-Whitney test (****p < 0.0001). Data from 76- 120 peroxisomes from n = 34-52 neurons and N = 6 independent experiments.

In axons, > 85% KIF21B-bound peroxisomes moved in the anterograde direction, consistent with the highly polarized nature of the axonal MT cytoskeleton (Figure 5E). KIF21B-bound peroxisomes in dendrites moved bi-directionally (Figure 5E), with 58% of the movement in the retrograde direction (Figure 5F). We compared the direction of motor recruited peroxisomes to the ∼ 60% plus-end out MT polarity measured in dendrites (Ayloo et al., 2017), and found that the 42% of KIF21B-FL recruited peroxisomes moved in the anterograde direction, which was significantly less than 60%, consistent with a retrograde bias. Strikingly, the average velocities of KIF21B-induced motion were significantly slower in dendrites than axons (Figure 5G), and the run distances were significantly shorter (Figure 5H). In addition, moving peroxisomes switched directions more frequently in dendrites than they did in axons (Figure 5I), consistent with the different MT arrangements in the two compartments. Thus, the 58% retrograde bias of KIF21B-induced motility observed in dendrites is the same as that observed by KIF21B puncta and KIF21B associated Trk/BNDF cargo in live hippocampal neurons (Ghiretti et al., 2016). In addition, this retrograde bias is remarkably similar to that observed by recruitment of the minus end-directed dynein-dynactin motor complex, which induced retrograde trafficking of 60% of peroxisomes (Ayloo et al., 2017), consistent with the net orientation of MTs.

### KIF21B recruited motors require both C-terminal MTRs and active MT dynamics to achieve a retrograde bias in dendrites

As our *in vitro* studies highlight the importance of the MBRs of KIF21B in regulating both motility and cytoskeletal remodeling, we asked if the MTRs are important for retrograde trafficking within dendrites of live neurons, and whether dynamic MT arrays are required for directionally biased motility. To determine if the MTRs impart retrograde bias within dendrites, we used our optogenetic recruitment assay to assess the direction and movement of full length (KIF21B-FL) and motor domain truncated (KIF21B-MD) motors when recruited to immobile peroxisomes in both axons and dendrites (Figure 5A). To investigate the role of MT dynamics, we used a low concentration of nocodazole (100 nM), which has been shown to dampen MT dynamics without net depolymerization (Guedes-Dias et al., 2019), in comparison to a DMSO control to test if regulation of MT dynamics is important for KIF21B retrograde bias. Recruitment of KIF21B motors to peroxisomes induced the processive motility of 53-76% of peroxisomes in axons and dendrites within 22-40 s, even in the presence of nocodazole (Figure S11A-F).

In axons, KIF21B-MD motors induced anterograde movement of > 95% of peroxisomes, similar to that seen with recruitment of KIF21B-FL motors (Figure 6A and C), even when MT dynamics were suppressed. However, in dendrites, removal of KIF21B’s C-terminal tail or addition of nocodazole led to anterograde movement of 53-57% of peroxisomes (Figure 6B and D), similar to that seen with recruitment of K560 motors (Ayloo et al., 2017). We did note substantial scatter in the data, likely due to heterogeneity in MT polarity along the length of dendrites, as shown in Figure 4 above.

**Figure 6.**
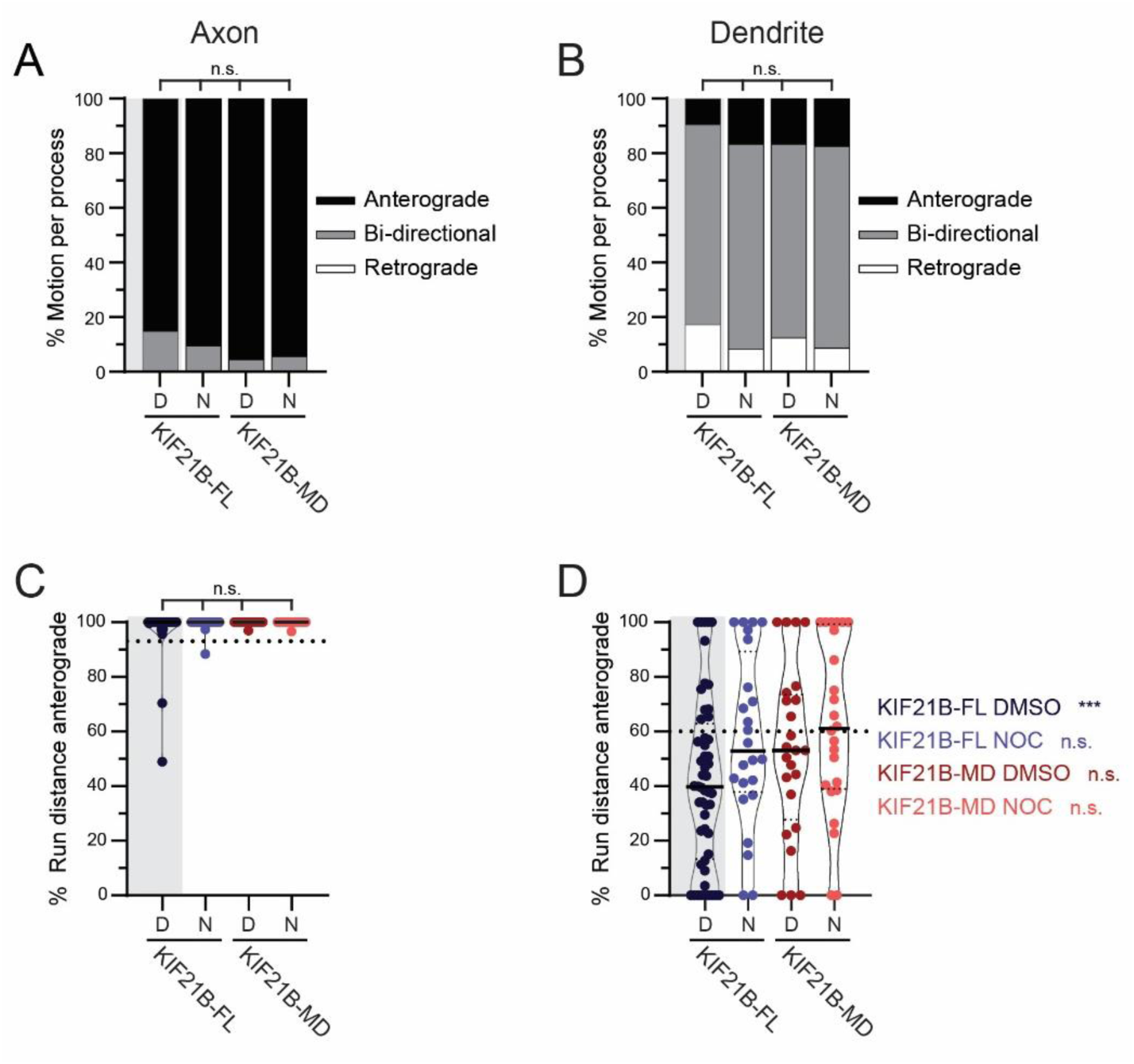
KIF21B induced retrograde trafficking in dendrites requires C-terminal tail domains and MT dynamics. A and B) Percentage of peroxisomes moving anterograde, retrograde or bi-directionally for individual axonal and dendritic processes. DMSO or nocodazole (N) conditions are indicated at the bottom of the graph. Bi-directional movement is characterized by peroxisomes that move more than 0.4 µm in both anterograde and retrograde directions. Kruskal–Wallis one-way ANOVA and Dunn’s multiple comparison (n.s. p > 0.05). C and D) Percentage of total peroxisome run distance per process moving in the anterograde direction in both axons and dendrites. DMSO (D) or nocodazole (N) conditions are indicated at the bottom of the graph. Plotted are individual values in color, means in solid black line and 25^th^ and 75^th^ quartiles in dashed black lines. For comparison of distributions we used Kruskal–Wallis one-way ANOVA and Dunn’s multiple comparison (n.s. p > 0.05). In dendrites we used a one sample Wilcoxon test to compare distributions against theoretical mean (60%). Significance is listed in color to the right side of the graph (n.s. p > 0.05; ***p < 0.001). Data from 37-120 peroxisomes from n = 18-52 neurons and N = 3-6 independent experiments.

We compared the direction of motor recruited peroxisomes in all conditions to the expected 60% plus-end out MT polarity measured in dendrites, which would induce 60% anterograde motility (Figure 6D). The direction of movement of KIF21B-FL recruited peroxisomes was significantly less than 60%, consistent with a retrograde bias. In contrast, 53% of KIF21B-MD recruited peroxisomes moved in the anterograde direction and did not show a similar retrograde bias. We also noted that when MT dynamics were dampened with nocodazole, 57% of KIF21B-FL and 62% of KIF21B-MD recruited peroxisomes moved in the anterograde direction, consistent with the 60% plus-end out directionality of MTs, suggesting that both KIF21B MTRs and MT dynamics are important for biasing movement in the retrograde direction.

There was no significant difference in the switching behavior of -FL or -MD recruited peroxisomes in either axons or dendrites (Figure S11A and B), even when MT dynamics were dampened. The frequency of directional switches seen by peroxisomes *in vivo* was greater than that seen *in vitro* on engineered MT bundles, suggesting that the recruitment of multiple KIF21B motor domains on cargo allows for robust track switching. In addition, recruitment of both KIF21B-FL and KIF21B-MD motors to peroxisomes induced similar velocities and run lengths in the presence and absence of nocodazole (Figure S11C-F). However, both KIF21B-FL and KIF21B-MD recruited cargos moved slower and less processively in dendrites than axons. Together, these results suggest that the C-terminal domains of KIF21B and MT dynamics are important for imparting retrograde biased movement in dendrites, but these factors do not dramatically affect the motility properties of KIF21B driven cargo transport.

## DISCUSSION

Most of our current motor trafficking knowledge comes from monitoring cargo movement within simple radial and axonal MT arrays (Burute & Kapitein, 2019; Hirokawa et al., 2010). Because of this, the motility properties of both kinesin and dynein are best understood in assays with unipolar MT arrays. However, there are many examples of cell types containing more complicated MT geometries on which molecular motors navigate (Muroyama & Lechler, 2017; Sanchez & Feldman, 2017). For example, kinesin and dynein motors function within the bipolar MT arrays of the mitotic spindle of dividing cells, where they work to pull and slide apart MTs, regulate MT dynamics, and transport materials and chromosomes to kinetochores and spindle poles (Dwivedi & Sharma, 2018; McIntosh, 2016; Wordeman, 2010). However, it is unclear how these motors establish directional movement along these antiparallel MT structures.

The mixed MT polarity characteristic of mammalian dendrites represents a similar challenge to understand how unidirectional motors can provide net long-range movement of cargos in a polarized manner. Kinesin motors have evolved specific properties to generate, maintain and navigate these complex arrays (Sweeney & Holzbaur, 2018). Some kinesin motors contain MT binding domains, in addition to their canonical motor domains, that have been implicated in sliding MTs against one another as well as transporting MTs throughout the cell (Andrews et al., 1993; Fink et al., 2009; Furuta & Toyoshima, 2008; Jolly et al., 2010; Navone et al., 1992; Reinemann et al., 2017; Seeger & Rice, 2010). In addition, many kinesin motors are known to influence the MT network by regulating MT dynamics and promoting MT assembly (Acharya et al., 2013; Arellano-Santoyo et al., 2017; G.-Y. Chen et al., 2019; Gudimchuk et al., 2013; Trofimova et al., 2018). Neuronal motors likely have evolved specific properties to control their directionality in dendrites.

Recent studies have advanced our understanding of the MT organization differences within dendrites and axons (Cao et al., 2020; Cunha-Ferreira et al., 2018; Liang et al., 2019; Sanchez & Feldman, 2017; Sánchez-Huertas et al., 2016; Wu & Akhmanova, 2017). Initial measurements using the “hook” method showed that axons contain uniform plus-end out oriented MTs, while only 52-57% of the MTs within dendrites are oriented with their plus-ends extending away from the cell body (Baas et al., 1988; Burton, 1988). Studies focused on dynamic MTs show that ∼ 65% of dendritic MTs are oriented plus-end out, as assessed by EB3 dynamics in live Purkinje neurons (Kleele et al., 2014; Stepanova et al., 2003). Previous measurements of EB3 comets in hippocampal neurons, at the same developmental stage as those used in this paper, indicated that 64% of MTs in dendrites are oriented plus-end out (Ayloo et al., 2017). The ∼ 60% anterograde motility that was observed for K560 motors moving on stabilized extracted dendritic MTs (Figure 4) is strikingly similar to the orientation of dynamic MTs within dendrites measured through EB3 comets (Ayloo et al., 2017). A model proposed by Kapitein and colleagues (Tas et al., 2017) suggests that the dendritic MT network is composed of distinct unipolar MT bundles, and that these bundles are marked by specific PTMs of the tubulin cytoskeleton. Using this same technique to track the movement of K560 motors, we acquired a detailed map of MT orientation within the dendrites of rat hippocampal neurons and found that MTs are organized heterogeneously into regions characterized by bundles with uniform polarity (Figure 3). The MT organization within axons and dendrites provides a clear way to differentiate between the two compartments (Bentley & Banker, 2016; Gumy & Hoogenraad, 2018; Nirschl et al., 2017). However, the mixed MT organization present within dendrites sets up problems for motors to move productively/efficiently within MT arrays.

KIF21B has been shown to move with net retrograde bias within the dendrites of live neurons while affecting MT dynamics (Ghiretti et al., 2016; Muhia et al., 2016). The *in vitro* assays reported here confirm that KIF21B strongly stabilizes MTs. We found that at physiologically relevant motor concentrations and ionic strength, KIF21B slows growth and potently suppresses catastrophe, leading to a net stabilization of the assembled MT (Figure1).

Using *in vitro* engineered MT bundles we observed that both motility and track switching by KIF21B were enhanced by MT bundling. Motor multimerization also had a robust effect on KIF21B run distance and track switching. KIF21B motors moving within native extracted dendrite MT arrays showed the directional switching seen along engineered MT bundles. Surprisingly, however, despite the pronounced track switching observed on extracted cytoskeletal arrays, KIF21B exhibited no directional bias in extracted dendritic MT arrays. In contrast, in live cells, acute recruitment of full-length KIF21B motors to non-motile peroxisomes using optogenetics was sufficient to induce both directional switching and active transport with a pronounced retrograde bias. These observations suggest that track switching may be necessary but not sufficient to establish a retrograde bias.

What controls KIF21B’s directional bias in dendrites? In live cells, KIF21B moved cargos in a net retrograde direction, but the motors showed no directional bias on extracted neuronal MT arrays. An important difference between the live cell and extracted network assays is the presence of dynamic MTs in live cells. When MT dynamics were dampened in dendrites of live neurons, the retrograde bias induced by KIF21B recruitment was eliminated. These results favor a model where modulation of MT dynamics by KIF21B is necessary for retrograde bias of cargo transport. In live neurons, KIF21B affects MT dynamics (Ghiretti et al., 2016; Muhia et al., 2016). In agreement, we observed that KIF21B robustly inhibits MT dynamics and assembly *in vitro*, which opens the possibility that this motor controls the remodeling of its own tracks. This remodeling requires the C-terminal domains of KIF21B containing the MTRs. Indeed, removal of the MTRs abolished retrograde biased cargo movement in live dendrites, suggesting that the MT dynamic regulation effect of the C-terminal MTRs is important for biasing retrograde biased movement. There is potential for this dynamic regulation to be different for more stable acetylated MTs oriented plus-end in over more dynamic tyrosinated MTs oriented plus-ed out due to differences in lattice stability (Janke & Magiera, 2020; Kelliher et al., 2019; Park & Roll-Mecak, 2018; Tas et al., 2017). In this case, KIF21B might differentiate between MT orientations by selectively walking on more stable plus-end in MTs, resulting in net retrograde bias. The synergistic combination of KIF21B-driven motility and modulation of MT dynamics may be necessary to direct KIF21B bound cargos in a net retrograde direction.

An alternate possibility is that the net retrograde bias is due to KIF21B reading the MT associated protein (MAP) code directly, rather than the PTM MT code. Regulatory MAPs could be lost during extraction, and dampening MT dynamics with nocodazole in live cell experiments could similarly affect MAP decoration. We did note that both KIF21B and K560 motor run distances were shorter on native and extracted dendritic MT arrays, compared to axonal arrays. This suggests that the native PTM MT code and MAP code, which are different in axons and dendrites, does affect long distance motor motility while having no effect on motor directionality.

In summary, our work provides evidence for a model where KIF21B motor teams utilize MT track switching and regulation of MT dynamics to traffic cargos with directional retrograde bias in the dendrites of neurons, despite the variable MT architecture (Figure 7). The ability of this motor to recognize and control the MT cytoskeleton sheds light on the morphological defects observed in both axon and dendrites where KIF21B is disrupted (Asselin et al., 2020; Morikawa et al., 2018; Muhia et al., 2016). Few kinesins have been directly implicated in dendritic transport, suggesting that specialized mechanisms are required to successfully navigate the complex MT organization within this cellular compartment. It will be of interest to understand if other dendritic kinesin motors navigate mixed MT arrays with a similar mechanism as that described here for KIF21B, or whether other motors have adopted alternative strategies.

**Figure 7.**
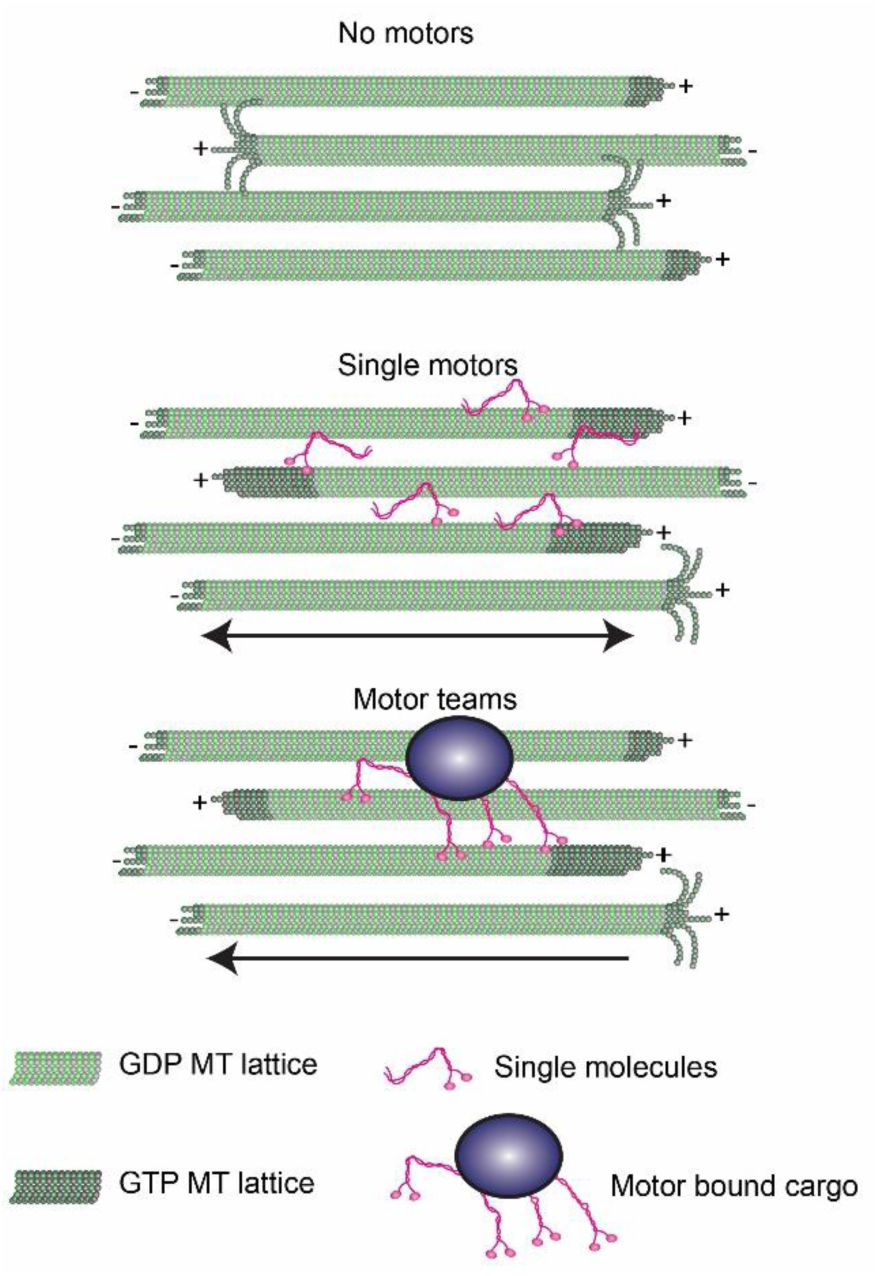
Model for KIF21B dendritic MT navigation. Single KIF21B motors contact MTs using N-terminal motor domains and MTRs in their C-terminal tails. Both single motors and motor teams contribute to stabilizing the dynamics of bound MTs. Binding of KIF21B motor domains and C-terminal MT binding sites stabilize the dynamics of bound MTs and increase the length of the GTP-tubulin cap at the plus ends. Motors also switch tracks between MTs closely positioned within arrays. The C-terminal tails of KIF21B promote track switching, and motor assembly along cargos allows for efficient track switching. In a region of mixed MT polarity, single motors move in both directions. However, KIF21B motors bound by cargos coordinate together to leverage track switching and regulation of MT dynamics to move long distances with net retrograde motility.

## MATERIALS AND METHODS

### Reagents

#### Constructs

For *in vitro* single molecule experiments, HaloTag-KIF21B donor plasmids were formed by inserting KIF21B sequences (full-length 1-1624 and motor domain 1-657) from prior constructs (Ghiretti et al., 2016) into pFastBac vectors. Baculovirus containing bacmid DNA for these constructs were produced at the Protein Expression Facility at The Wistar Institute.

The K560-GFP plasmid (Addgene #15219) was acquired from the Vale lab (Woehlke et al., 1997). The K560 sequence was inserted into the pHTC HaloTag CMV neo vector (Promega, Madison, WI, US, G7711) to build a K560-Halo plasmid. The rigor-kinesin (K560 E236A mutant, Addgene #60909) plasmid was acquired from the Vale lab (Tanenbaum et al., 2014)

For peroxisome recruitment, the KIF21B-mCherry-eDHFR construct was derived by inserting the KIF21B sequence from the mCherry-KIF21B construct (gift of Matthias Kneussel, University of Hamburg) into the mCherry-eDHFR plasmid previously described in Ayloo et al., (2017). The GFP-Halo-Pex3 construct was used as described in Ayloo et al., (2017).

#### Protein Expression and Purification

To express KIF21B-FL-Halo and -MD-Halo proteins, Sf9 insect cells (Expression Systems, Davis, CA, US, 94-001F) were grown to a density of 4x106 cells/mL in ESP 921 media (Expression Systems, 96- 001-01) at 27 °C, at which point they were infected with high titer baculovirus for 48 hr. Cells were pelleted and flash frozen in liquid nitrogen and stored at −80 °C. On the day of purification, cells were lysed in lysis buffer (10 mM Tris, 200 mM NaCl, 2 mM ATP, 4 mM MgCl_2_, 5 mM DTT, 1 mM EGTA, 0.5% Igepal, 1 mM β-ME, and 0.01 mg/mL aprotinin (GoldBio, St. Louis, MO, US, A-655-100) and leupeptin (Peptides International, Louisville, KY, US, ILP-4041-100mg), pH 7.5) and clarified through centrifugation at 42,000 rpm for 1 hr. Lysate was flowed over an Anti-FLAG resin (GenScript, Piscataway, NJ, US, L00432) column at 1 mL/min. After washing bound protein with 10 column volumes (CVs) wash buffer (10 mM Tris, 200 mM NaCl, 2mM ATP, 4 mM MgCl_2_, 5 mM DTT, 1 mM EGTA, 1 mM β-ME, and 0.01 mg/mL aprotinin and leupeptin, pH 7.5), protein was eluted by incubating 1 CV of elution buffer (10 mM Tris, 200 mM NaCl, 1 mM DTT, 1 mM EGTA, 0.2 mg/mL FLAG peptide (Sigma-Aldrich, St. Louis, MO, US, SAB1306078-400UL), pH 8.0) with resin beads for 1 hr, followed by a second 1 CV incubation for 30 min, and three more quick 1 CV incubations for a total of 5 CVs elution volume. Eluted protein was labeled with either 1.75 µM AlexaFluor 660 or 2.5 µM TMR HaloTag Ligands (Promega, G8471 and G8251) for 2 hr on ice. Labeled protein was dialyzed in PEM buffer (100 mM NaPIPES, 50 mM NaCl, 1 mM MgSO_4_, 1 mM EGTA, pH 6.8) for 2-18 hr. Protein was then flowed over a Q sephaorse (GE Healthcare, Chicago, IL, US, 17051001) column, and eluted in high salt buffer (100 mM NaPIPES, 600 mM NaCl, 1 mM MgSO_4_, 1 mM EGTA, pH 6.8). MT affinity/dead-head spin was performed by binding motor protein to newly polymerized MTs (unlabeled, 2uM Paclitaxel (Taxol (Cytoskeleton, Denver, CO, US, TXD01) -stabilized) in binding buffer (12 mM NaPIPES, 200 mM KCl, 5 mM MgCl_2_, 2 mM EGTA, 1 mM AMPPNP, and 20 µM Taxol, pH 6.8) for 30 min. Motors and MTs were pelleted through centrifugation at 18 krpm for 20 min. Un-bound motors were rinsed using wash buffer (12 mM NaPIPES, 200 mM KCl, 5 mM MgCl_2_, 2 mM EGTA, and 20 µM Taxol, pH 6.8). Bound motors were released from MTs through a 5 min incubation in release buffer (12 mM NaPIPES, 200 mM KCl, 5 mM MgCl_2_, 2 mM EGTA, 5 mM ATP, and 20 µM Taxol, pH 6.8), followed by centrifugation at 18 krpm for 10 min to remove any MTs and non-active kinesin motors from the supernatant. MT-released protein was aliquoted, flash frozen and stored in liquid nitrogen.

To produce K560-Halo and K560-GFP protein, plasmids were transformed into BL21(DE3)pLysE bacteria (Sigma-Aldrich, CMC0015-20X40UL) and grown in TB supplemented with 50 µg/mL kanamycin and 33 µg/mL chloramphenicol at 37 °C until an OD_600_ of 0.4 was reached. Cultures were cooled to 18 °C and protein expression was induced with 0.15 mM IPTG for 18 hr. Cells were pelleted and flash frozen in liquid nitrogen and stored at −80 °C. On the day of purification, cells were lysed by microfluidizer (Microfluidics, Westwood, MA, US) after resuspension in a lysis buffer (50 mM NaPO_4_, 250 mM NaCl, 20 mM Imidazole, 1 mM MgCl_2_, 0.5mM ATP, 1 mM β-ME, 0.01 mg/mL aprotinin and leupeptin, pH 6.0), and clarified through centrifugation at 42,000 rpm for 30 min. Lysate was run over a Co2+ agarose bead (GoldBio, H-310-25) column at 1 mL/min. After washing bound protein with 10 CVs wash buffer (50 mM

NaPO_4_, 300 mM NaCl, 10 mM Imidazole, 1 mM MgCl_2_, 0.1 mM ATP, 0.01 mg/mL aprotinin and leupeptin, pH 7.4), protein was eluted with 5 x 1 CV of elution buffer (50 mM NaPO_4_, 300 mM NaCl, 150 mM Imidazole, 1 mM MgCl_2_, 0.1 mM ATP, pH 7.4). Fractions were pooled and concentrated. Buffer was exchanged to BRB80 (80 mM NaPIPES, 1 mM MgCl_2_, 1 mM EGTA, pH 6.8) by loading protein over NAP- 10 (GE Healthcare, 17-0854-01) and PD-10 (GE Healthcare, 17-0851-01) desalting columns. MT affinity/dead-head spin was performed as described above for KIF21B motors. MT-released protein was aliquoted, flash frozen and stored in liquid nitrogen.

To produce rigor-kinesin protein, the construct plasmid was transformed into BL21(DE3)pLysE bacteria (Sigma-Aldrich, CMC0015-20X40UL), and grown in TB supplemented with 100 µg/mL ampicillin and 33 µg/mL chloramphenicol at 37 °C until an OD_600_ of 0.4 was reached. Cultures were cooled to 18°C and protein expression was induced with 0.1 mM IPTG for 18 hr. Cells were pelleted and flash frozen in liquid nitrogen and stored at −80 °C. On the day of purification, cells were lysed by microfluidizer after resuspension in a lysis buffer (25 mM KPO_4_, 300 mM NaCl, 40 mM Imidazole, 1 mM MgCl_2_, 10% glycerol, 100 µM ATP, 670 mM PMSF, pH 8.0), and clarified through centrifugation at 42,000 rpm for 30 min. Lysate was run over a Co2+ agarose bead column at 1 mL/min. After washing bound protein with 10 CVs of wash buffer at pH 8.0 (300 mM NaCl, 40 mM Imidazole, 1 mM MgCl_2_, 10% glycerol, 100 µM ATP, 5 mM β-ME, pH 8.0), and 10 CVs wash buffer at pH 7.0, protein was eluted with 5 CVs of elution buffer (300 mM NaCl, 250 mM Imidazole, 1 mM MgCl_2_, 10% glycerol, 100 µM ATP, pH 7.0). Elution fractions were pooled and concentrated. Protein was aliquoted, flash frozen and stored at -80 °C.

#### Neuronal Cell Culture

The day before plating neurons, 35-mm glass-bottom dishes (MatTek, Ashland, MA, US, P35G- 1.5-14-C) or 25 mm round coverslips were coated with 0.5 mg/mL poly-L-lysine (Sigma-Aldrich, P1274). E18 Sprague–Dawley rat hippocampal neurons were received from the Neuron Culture Service Center at the University of Pennsylvania and plated in attachment media (MEM (Gibco®, ThermoFischer, Waltham, MA, US, 1109-072) supplemented with 10% horse serum (Gibco®, 26050-070), 33 mM glucose (Corning, Corning, NY, US, 25-037-CIR), and 1 mM sodium pyruvate (Gibco®, 11360-070)) at a density of 100,000 cells per coverslip or 500,000 cells per dish. After 4-6 hr of attachment, neurons were cultured at 37 °C with 5% CO_2_ and maintained in either maintenance media (Neurobasal (Gibco®, 21103-049) supplemented with 33 mM glucose (Corning, 25-037-CIR), 2 mM GlutaMAX (Gibco®, 35050-061), 100 units/mL penicillin and 100 µg/mL streptomycin (Gibco®, 15140-122), and 2% B27 (Gibco®, 17504-044)) or BrainPhys Neuronal Medium with NeuroCult SM1 Neuronal Supplement (BrainPhys Primary Neuron Kit (StemCell Technologies, Vancouver, CA, 05794)). AraC (Sigma-Aldrich, C6645) was added at 10 µM 24 hr after initial plating to prevent glial cell division.

#### Antibodies

Antibodies and dilutions used in immunofluorescence assays were: rat anti-Tyrosinated α- Tubulin, clone YL1/2 (EMD Millipore Corporation, Billerica, MA, US, MAB1864, 1:500), mouse anti-Acetylated Tubulin, clone 6-11B-1 ((Sigma-Aldrich, T7451, 1:1000), goat anti-Rat IgG (H+L) Alexa Fluor 488 (ThermoFischer, A-11006, 1:1000), and goat anti-Mouse IgG (H+L) Alexa Fluor 555 (ThermoFischer, A-21424, 1:1000).

### Experimental Procedures

#### *In Vitro* Single Molecule Assay

Flow cells were assembled from attaching silane (PlusOne Repel Silane, GE Healthcare, 17- 1332-01) coated coverslips and cleaned glass slides together with double-side stick tape to form ∼ 10 μL flow chambers. Each flow cell was treated as follows: (1) 10 μL 1µM rigor-kinesin incubated for 5 min; (2) 30 μL 5% pluronic F-127 (Sigma-Aldrich, P2443-250G) incubated for 5 min; (3) 30 μL casein wash buffer (30 mM DTT, 1 mg/mL filtered casein (Sigma-Aldrich, C5890-500G) in BRB80 pH 6.8); (4) 50 μL of 1 μM MT seeds incubated for 2 min. Seeds were formed by shearing larger MTs (GMPCPP-stabilized, labeled 1:40 with HiLyte 488 tubulin (Cytoskeleton, TL488M-B) with a Hamilton syringe (Avanti Polar Lipids, Alabaster, AL, US, 610015). Seeds were aligned in the direction of flow by flowing in the seed solution with the chamber glass at a low grade angle to keep the flow slow and continuous; (5) 30 μL casein wash buffer; (6) 20 μL final flow mixture incubated while imaging. Final flow: 0-50 nM kinesin motor, 10 µM tubulin (labeled 1:20 with HiLyte 488 tubulin, 1 mM ATP, 1 mM GTP, 0.32 mg/mL casein, 0.32 mg/mL bovine serum albumin (BSA) (Fischer Scientific, ThermoFischer Scientific, 50-253-90), 9.7 mM DTT, 15.5 glucose, 119 U/mL glucose oxidase (Sigma-Aldrich, G2133-250KU), 278 U/mg catalase (Sigma-Aldrich, C100-500MG), 0.244 mg/mL creatine phosphokinase (Sigma-Aldrich, C3755-35KU), 11.5 mM phosphocreatine (Sigma-Aldrich, P7936-1G, and 0.2% Methylcellulose, diluted in BRB80 pH 6.8. For KIF21B-FL multimerization experiments, final flow was diluted in P12T (12 mM NaPIPES, 2 mM MgCl_2_, 1 mM EGTA, pH 6.8).

#### TIRF Nucleation Assay

Flow cells were assembled by attaching silane coated coverslips and cleaned glass slides together with double-side stick tape to form ∼ 10 μL flow chambers. Each flow cell was blocked with 5% pluronic F-127 incubated for 5 min and washed with casein wash buffer (30 mM DTT, 1 mg/mL filtered casein in BRB80 pH 6.8) prior to addition of final flow solution. A final flow (0-50 nM kinesin motor, 10 µM tubulin (labeled 1:20 with HiLyte 488 tubulin, 1 mM ATP, 1 mM GTP, 0.32 mg/mL casein, 0.32 mg/mL BSA, 9.7 mM DTT, 15.5 glucose, 119 U/mL glucose oxidase, 278 U/mg catalase, 0.244 mg/mL creatine phosphokinase, 11.5 mM phosphocreatine, and 0.2% Methylcellulose, diluted in BRB80 pH 6.8) was added to chambers and heated to 35°C with an objective collar. Chambers were imaged 10 min after incubation with final flow solution.

#### Light Scattering Assay

Wells of a 96-well half area UV transparent plate (Corning, 3679) were used to mix solutions containing 50 nM kinesin motor, 10 µM Taxol or BRB80 in assay buffer (10 µM tubulin, 10% glycerol (v/v), 4 mM DTT, 1 mM GTP, 1 mM ATP, and 0.133 mg/mL casein diluted in BRB80 pH 6.8). Solutions were pipetted as triplicates. Absorbance values of each well were measured at 340 nm every min at 37 °C using a prewarmed SynergyMx platereader (BioTek).

#### Motor-PAINT Assay

MT arrays in neurons were fixed and stabilized as described (Tas et al., 2017) with slight modifications. Briefly, membranes were removed from neurons cultured in MatTek dishes at 8-10 days in vitro (DIV) by incubating cells with extraction buffer (1M sucrose, 0.15% Triton X-100 (Roche, Basel, CH, MilliporeSigma, 10789704001) in BRB80 pH 6.8 at 37 °C) for 1 min. An equal amount of fixation buffer (1% paraformaldehyde (PFA (Affymetrix, ThermoFischer Scientific, 199431LT) in BRB80 pH 6.8 at 37 °C) was added for 1 min with gentle swirling. Dishes were rinsed 3 times with wash buffer (2 µM Taxol in BRB80 pH 6.8 at 37°C). Dishes were washed once more right before imaging. To image, wash buffer was replaced with imaging buffer (1 mM ATP, 2 µM Taxol, 0.133 mg/mL casein, 0.133 mg/mL BSA, 4 mM DTT, 6 mg/mL glucose, 49 U/mL glucose oxidase, 115 U/mg catalase, 0.21 mg/mL creatine phosphokinase, and 4.76 mM phosphocreatine in BRB80 pH 6.8) containing 5-10 nM KIF21B-Halo-660 motor and 5-10 nM K560-GFP motor.

#### Immunofluoresence

At 8-10 DIV, hippocampal neurons plated coverslips were either extracted and fixed as described for motor-PAINT assay or alternatively fixed with 37 °C warmed 4% PFA in PBS (50 mM NaPO_4_, 150 mM NaCl pH 7.4) for 10 min, followed by 3 PBS washes and permeabilization in 0.1% Triton-X100 in PBS for 10 min. Coverslips were blocked with cell block (0.2% Triton-X100 and 3% BSA in PBS) for 2 hr at room temperature. Primary antibodies were diluted as described above in cell block and incubated on coverslips for 2 hr at room temperature. Coverslips were then washed three times, for 10 min each, with PBS. Alexa-conjugated secondary antibodies were diluted in cell block and incubated on coverslips for 1 hr at room temperature. Coverslips were washed again three times with PBS, each for 10 min. Coverslips were rinsed in dH_2_O and mounted on glass slides with ProLong Gold antifade reagent (Invitrogen, P36930).

#### Induced Recruitment Assay

Induced recruitment experiments with KIF21B motors was performed in DIV 8-10 hippocampal neurons as previously described (Ayloo et al., 2017), with the exception of transfecting hippocampal neurons with DNA plasmids for PEX3-GFP-Halo and either full-length KIF21B(aa1-1624)-mCherry-eDHFR or motor domain truncated KIF21B(aa1-657)-mCherry-eDHFR. Cells were incubated with 100 nM Nocodazole (Sigma-Aldrich, M1404-10MG), or dimethyl solfoxide (DMSO, ACROS Organics, Geel, BE, 326881000) as well as 10 µM cTMP-Htag (Ballister et al., 2015) or CTH (Zhang et al., 2017) dimerizer 30 min before imaging.

#### Microscopy

Single molecule and Motor-PAINT assays with dynamic MTs were performed at 37 °C using a PerkinElmer Nikon Eclipse Ti TIRF system, using a Nikon Apo TIRF 100x 1.49 NA oil-immersion objective and a Hamamatsu ImagEM C9100-13 EMCCD camera operated by Volocity software (PerkinElmer, Waltham, MA, US, version 6.4.0). TIRF MT nucleation experiments were performed at 35 °C using a dual-view Leica TIRF microscope, with an Olympus UplanApo 60x 1.45 NA oil-immersion objective and an Andor iXon Ultra EMCCD camera operated with Metamorph software (Molecular Devices, San Jose, CA, US, version 7.10.3.279). Immunofluorescence images were acquired with a Leica DMI6000 inverted microscope, using an HCX PL APO 40x 1.25 NA oil-immersion objective and a Hamamatsu ORCA-R2 CCD camera operated with Leica Application Suite X (LAS X) software (Leica Microsystems, Buffalo Grove, IL, US, version 3.0.3.279). Induced recruitment assays were performed at 37 °C using a Perkin Elmer Nikon Eclipse spinning-disk confocal Ultraview VoX system equipped with a 405 nm Ultraview Photokinesis accessory, using a Hamamatsu ImagEM C9100-50 EMCD camera operated by Volocity software.

Movies in each experiment were taken with different frame rates. Motor-PAINT movies were obtained by continuously acquiring motor images at 5 frames/s (FPS) for 2 min. Peroxisome-motor induced recruitment movies were taken by imaging both peroxisome and motor images at 2 FPS for 20 s prior to photoactivation, followed by 2 FPS for 2-5 min after photoactivation. *In vitro* assay bleaching movies of KIF21B multimers was acquired by continuously acquiring motor images at 5 FPS for 5-15 min. *In vitro* assay movies of KIF21B multimers and single molecules were obtained by acquiring motor images at 5 FPS while acquiring MT track images at 1 frame/second for 2 min. Every 6^th^ frame of these movies was skipped to image the growing MTs. *In vitro* assays of dynamic MTs were obtained by continuously acquiring MT images at 2 FPS for 10 min.

### Quantification and Statistical Analysis

#### Protein composition

Recombinant protein concentration was assessed using a Pierce BCA Protein Assay Kit (ThermoFischer, 23225). Protein purity was assessed through western blotting with Odyssey CLx Imaging System (LI-COR Biosciences, Lincoln, NE, US) using Image Studio Lite version 5.2.5.

#### Motor Multimerization Calibration

We measured the number of TMR Halo tag ligands bound to each KIF21B peptide through stepwise bleaching as well as the corresponding intensity values for each step. We then fit a linear equation to the data to describe the number of TMR dyes and the measured intensity and calculated the intensity for 2 TMR dyes, corresponding to one motor dimer to within one standard deviation. We applied the ratio of this value to the mean intensity of the population of all spots measured to that of the moving spot data to determine the intensity threshold limit of a motor dimer.

#### Single Particle Tracking

Single kinesin particles moving along extracted axonal and dendritic MT arrays were segmented out with Cega (Masucci, Relich et al., 2020) and tracked with a modified version of the software used in Schwartz et al., (2017) and described in Relich, (2016), which is a tracking algorithm inspired by u-track (Jaqaman et al., 2008). Cega modified the initial candidate motor finding step to greatly enhance the quality of the downstream tracking processes when analyzing data that has heterogeneous background fluorescence and nuisance particles. To eliminate particles moving along arrays belonging to another nearby process, a dilated binary mask was applied. Once the candidates were identified, the original movie images were analyzed by the localization algorithm with tracking performed by an algorithm similar to Schwartz et al., (2017). Analysis of track segment directionality was performed for segments that moved within 50-400 nm and belonged to a track with more than 3 connected segments. Intensity information for track segments was calculated from the original unmodified movie data.

#### Image/Kymograph Analysis

For single molecule experiments using dynamic MTs, growth and catastrophe events along growing MTs were characterized by a change in the MT plus-end position that exceeded 0.5 µm. Dynamic MT tracing was accomplished using MATLAB R2019a software (MathWorks, Natick, MA, US, version 9.6.0.1174912). Purified KIF21B motility and dynamic MT behavior was analyzed with kymographs generated along the MT long-axis. Unless specifically states, we focused on analyzing only motor particles whose intensity was consistent with single molecules, of which of which ∼ 50% were fully labeled. Run distances and velocities were quantified for motor runs that extend 0.5 µm (3 pixels) or more. Run distances were measured as the sum of run lengths in each direction for an individual motor. Track switches are characterized by a reversal in run direction that exceeds 0.5 µm. Along extracted MT arrays the direction switches for each kinesin was quantified by analyzing kymographs of motor movement along the midline of each process. The frequency of switching was calculated by counting the number of motors that switched direction or did not switch direction along the center line of each kymograph. Switches were quantified for motor runs that extended 0.5 µm or more.

For neuron experiments, axons and dendrites were identified based on morphologic criteria as previously outlined (Kaech & Banker, 2006). At 8-10 DIV, dendrite lengths in our cultures were ∼ 50-100 µm. For motor induced recruitment experiments, only neurons expressing both of the co-transfected GFP and mCherry markers were imaged. Peroxisome motility was analyzed using kymographs generated along the length of each process using the Multi Kymograph plugin for Fiji version 1.8.0-66 (ImageJ 1.53c, National Institutes of Health, Bathesda, MD). Peroxisome diameter was measured along the long axis of each corresponding process. Run distances and velocities were quantified for peroxisomes that moved greater than 3 µm. Run distances were measured as the total sum of distances run in each direction for an individual organelle. Peroxisome movement was considered bi-directional if the organelle moved greater than 0.4 µm in both the anterograde and retrograde directions. Peroxisome switches were quantified for organelles that moved more than 0.4 µm in the opposing direction.

#### Statistical Analysis

Data from each experiment was analyzed from a minimum of three independent replicates, unless otherwise noted. Our statistical analysis was performed in GraphPad Prism (version 9.0.1 (151), Prism 9, San Diego, CA, US). We used a two-tailed Mann-Whitney test to compare two variables and Kruskal-Wallis ANOVA with Dunn’s multiple comparisons or Ordinary one-way ANOVA to compare multiple variables. A Wilcoxon signed-rank test was used to determine if values were significantly different from 60%, and an F test was used to determine if values were significantly different from zero. Statistical values are mentioned in figure legends.

### DATA AVAILABILITY

Source data files will be provided for all the Figures. Data files will be available at doi:10.5281/zenodo.4558240.

## Supporting information

Video 1. Purified KIF21B-FL motor motility along single dynamic MTs.

Video 2. Purified KIF21B-FL motor motility along dynamic MTs forming a parallel bundle.

Video 3. Purified KIF21B-FL motor motility along dynamic MTs forming an antiparallel bundle.

Video 4. Purified K560-GFP motor motility along extracted axonal MT arrays.

Video 5. Purified K560-GFP motor motility along extracted dendritic MT arrays.

Video 6. Induced recruitment of KIF21B motors to peroxisome cargo in axons of live neurons.

Video 7. Induced recruitment of KIF21B motors to peroxisome cargo in dendrites of live neurons.

## ACKNOWLEDGMENTS

We acknowledge Mariko Tokito for help designing and constructing DNA constructs; Daniel Safer for help producing recombinant proteins; and Amy Ghiretti for help optimizing and performing induced recruitment experiments. This work was supported by the Center for Engineering MechanoBiology NSF Science and Technology Center, CMMI:15-48571, and National Institutes of Health grants RM1 GM136511 (to E.M.O, E.L.F.H and M.L.) and R35 GM126950 (to E.L.F.H).

## SUPPLEMENTAL FIGURES

**Figure S1.**
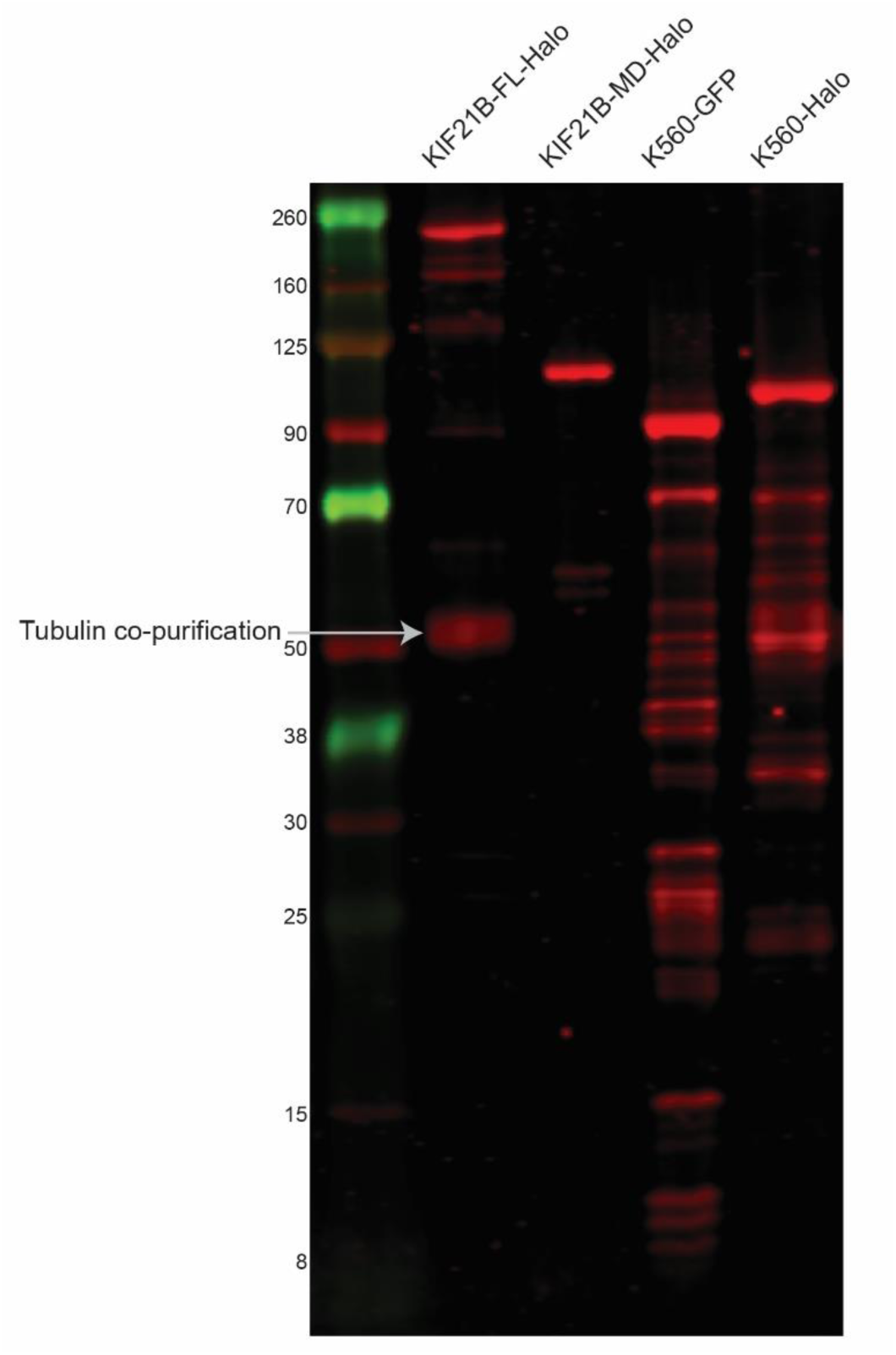
Western blot analysis of recombinant proteins. Revert 700 Total Protein Stain of purified kinesin proteins. 10 μL of 1-2 µM protein was loaded per lane.

**Figure S2.**
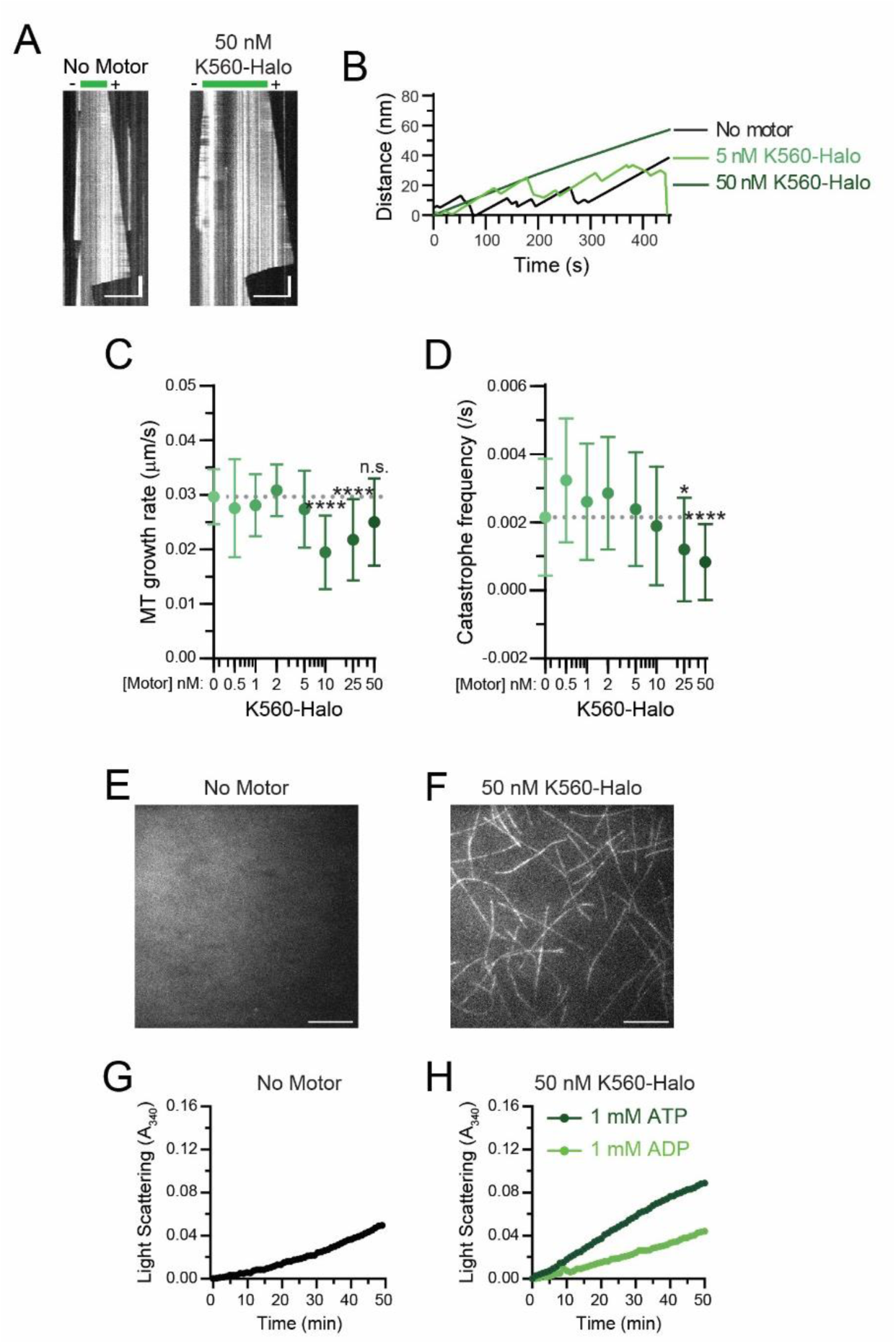
K560 motors stabilize MT dynamics and promote MT assembly. A) Kymographs showing dynamic MTs polymerizing from stabilized MT seeds in the absence or presence of K560-Halo. Plot for the no motor condition was repeated from figure 1 for comparison. Scale bars: 10 µm (horizontal) and 30 s (vertical). B) Average MT plus-end growth trace. For each condition 10 MT plus-end trajectories were averaged. C) MT plus-end growth speed in the presence of increasing concentrations of K560-Halo. Plotted are means and standard deviations. Kruskal–Wallis one-way ANOVA and Dunn’s multiple comparison (n.s. p > 0.05; ****p < 0.0001). D) MT plus-end catastrophe frequency in the presence of increasing concentrations of K560-GFP. Plotted are mean and standard deviations. Kruskal–Wallis one-way ANOVA and Dunn’s multiple comparison (*p < 0.05; ****p < 0.0001). E-F) TIRF microscopy images of free tubulin dimers incubated in the absence or presence of 50 nM K560-Halo. Images were taken 10 min after solutions were added and incubated in flow chambers. Scale bar: 10 µm. G-H) Light scattering traces for solutions containing buffer, MT polymerizing and depolymerizing drugs, 50 nM K560-Halo with 1 mM ATP or 1 mM ADP. Graphs show mean values for each time point (See Figure S3 for graphs with standard deviations). Means were compared with Kruskal–Wallis one-way ANOVA and Dunn’s multiple comparison. (Buffer – K560-Halo ****p < 0.0001; K560-Halo ATP – K560-Halo ADP ****p < 0.0001). Data from 7-18 traces from N = 5 independent experiments. Unless otherwise indicated, data from 51-74 MTs and N = 4-5 independent experiments.

**Figure S3.**
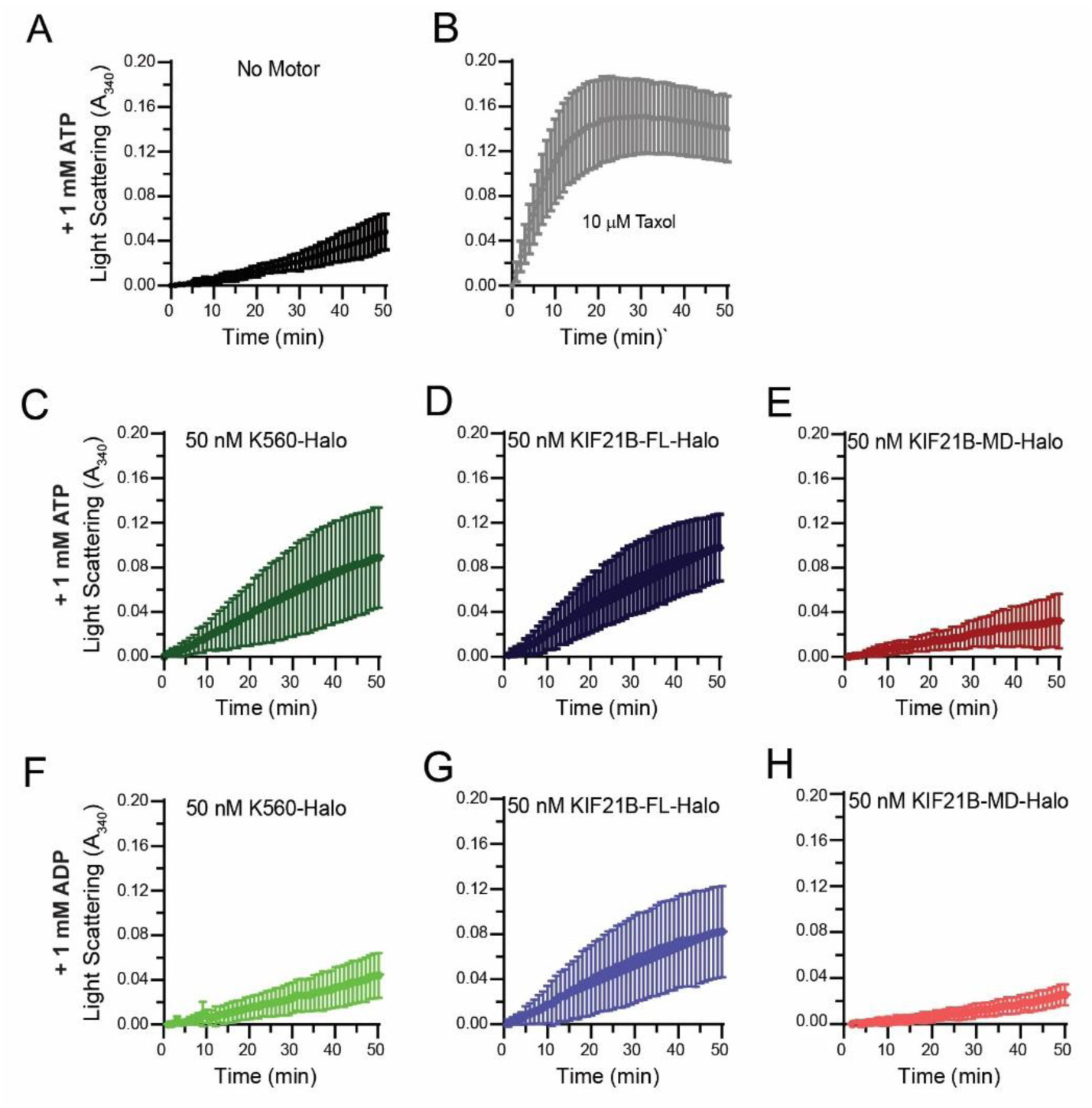
KIF21B C-terminal MTRs promote MT assembly. Light scattering traces for solutions containing buffer, MT polymerizing and depolymerizing drugs, or kinesin motors with 1 mM ATP or 1 mM ADP. Graphs show mean values and standard deviations for each time point. Means were compared with Kruskal–Wallis one-way ANOVA and Dunn’s multiple comparison. Data from 7-22 traces from N = 5 independent experiments.

**Figure S4.**
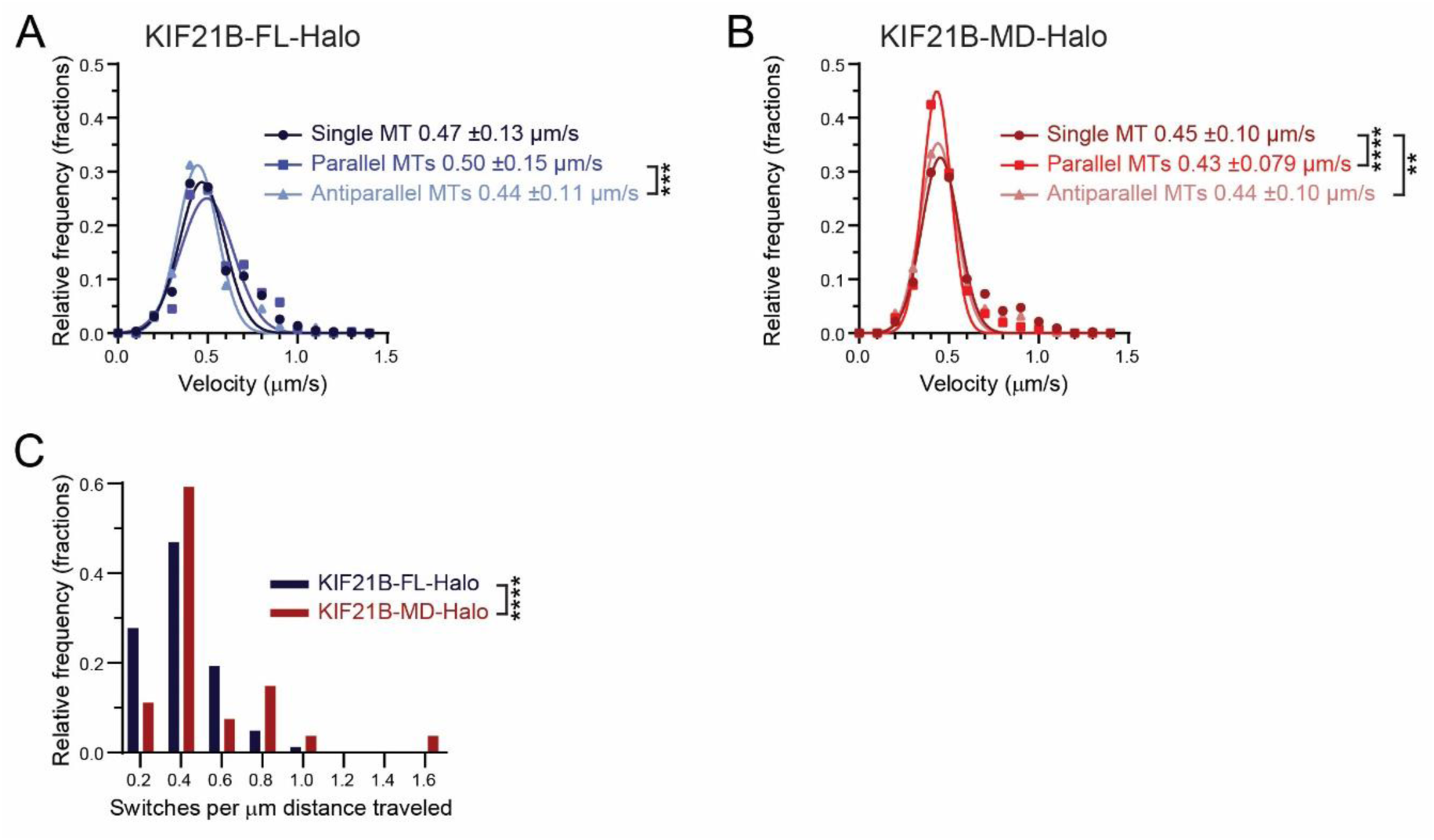
KIF21B MTBs promote track switching, but do not affect motor velocity. A and B) Histogram of velocities of single KIF21B-FL-Halo and KIF21B-MD-Halo molecules along single MTs or MT bundles. Data points were fitted by a gaussian function. Listed are the means and standard deviations. Ordinary one-way ANOVA (**p < 0.01; ***p < 0.001; ****p < 0.0001). C) Histogram of the number of track switches per distance traveled along antiparallel MTs for KIF21B-FL-Halo and KIF21B-MD-Halo motors. Two-tailed Mann-Whitney (****p < 0.0001). Data from 400-1069 motor runs from n = 20-60 MTs and N = 6-7 independent experiments.

**Figure S5.**
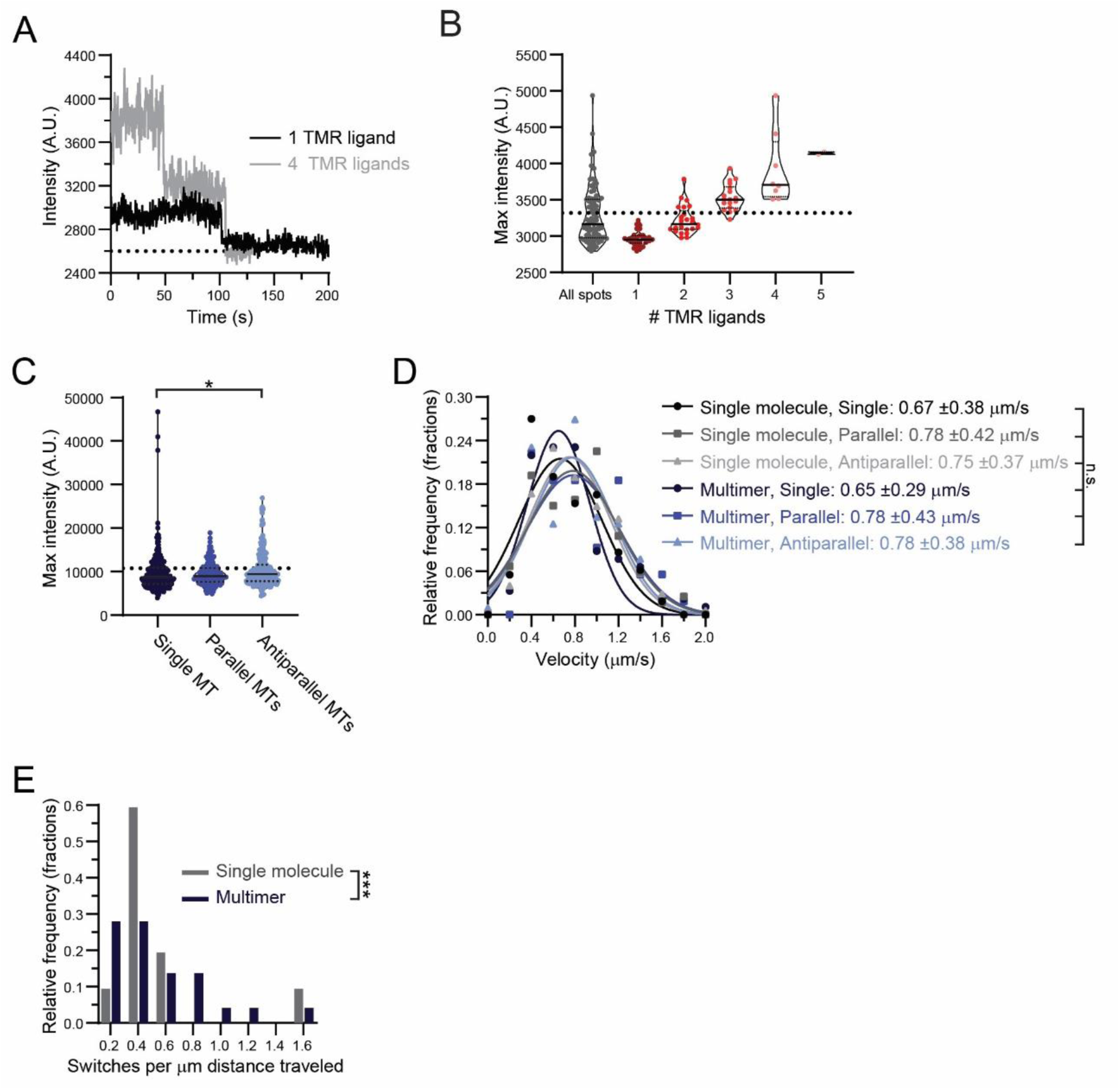
Calibration of single molecule and multimer intensity threshold and velocity quantification for purified KIF21B motors in low ionic strength conditions. A) Example intensity profiles for TMR Halo ligand quenching on KIF21B-FL-Halo motors. Black traces indicate quenching of one TMR dye and gray traces indicate quenching of four TMR molecules. Dashed black line represents average background intensity (2,632 A.U.). B) Maximum intensity distribution of KIF21B spots versus distributions representing increasing TMR dye number. The threshold separation between 2 and 3 TMR molecules is indicated with the black dashed line (3,343 A.U.). Plotted are individual values in color and means in solid black lines and 25^th^ and 75^th^ quartiles in dotted black lines. Data from 2-83 motor runs from n = 20 MTs and N = 1 independent experiment. C) Maximum intensity distribution of KIF21B particles moving on different MT types. The threshold for single molecule intensity is indicated with the black dashed line (10,117 A.U.). Plotted are individual values in color and means in solid black lines and 25^th^ and 75^th^ quartiles in dotted black lines. Ordinary one-way ANOVA (*p < 0.05). Data from 54-174 motor runs from n = 11-12 MTs and N = 4 independent experiments, unless otherwise stated. D) Histogram of velocities of single molecule and multimeric KIF21B-FL-Halo motors along single MTs or MT bundles. Data points were fitted by a gaussian function. Listed are the means and standard deviations. Ordinary one-way ANOVA (n.s. > 0.05). E) Histogram of the number of track switches per distance traveled along antiparallel MTs for single molecule and multimeric KIF21B-FL-Halo motors. Two-tailed Mann-Whitney (***p < 0.001). Unless otherwise stated, data for single and multimeric KIF21B-FL-Halo from 54-174 motor runs from n = 11-12 MTs and N = 4 independent experiments.

**Figure S6.**
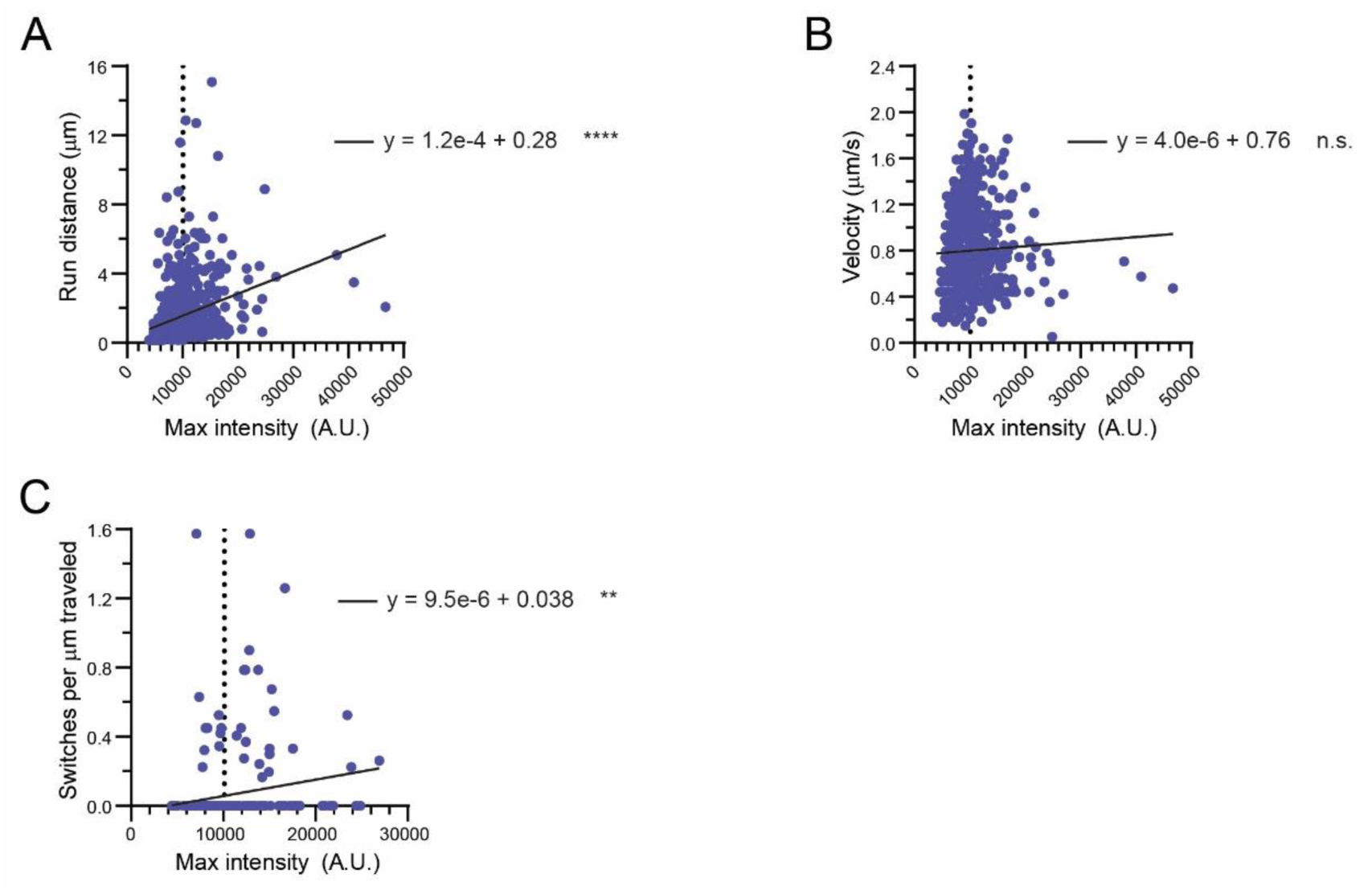
KIF21B motor intensity is positively correlated with run distance and track switching, and not correlated with velocity. A) Scatter plot of maximum motor intensity versus run distance. Plotted is a linear regression line for KIF21B-FL-Halo motors. Black dashed line indicates intensity threshold for single molecules. F test (****p < 0.0001). B) Scatter plot of maximum motor intensity versus velocity. Plotted is a linear regression line for KIF21B-FL-Halo motor spots. Black dashed line indicates intensity threshold for single molecules. F test (n.s. p > 0.05). C) Scatter plot of maximum motor intensity versus switches per run distance. Plotted is a linear regression line for KIF21B-FL-Halo spots. Black dashed line indicates intensity threshold for single molecules. F test (** p < 0.01). Data from 54-174 motor runs from n = 11-12 MTs and N = 4 independent experiments.

**Figure S7.**
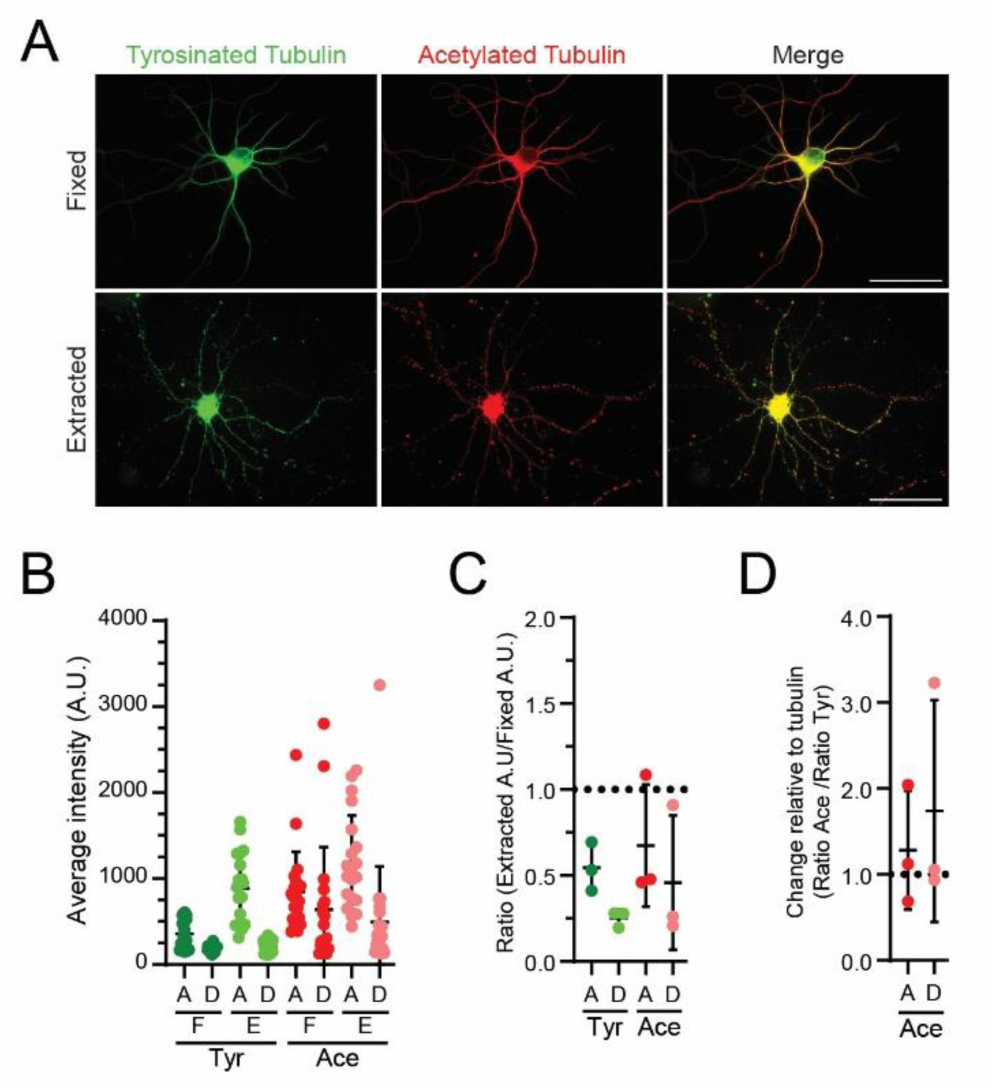
Tubulin PTMs are maintained on extracted MT arrays from cultured hippocampal neurons. A) Representative widefield images of fixed or extracted hippocampal neurons stained for tyrosinated and acetylated tubulin. Brightness and contrast levels are adjusted differently for each image. Scale bars: 50 µm. B) Average intensity values for secondary antibodies indicating tyrosinated (Tyr) and acetylated (Ace) tubulin. Values were averaged along axons (A) and dendrites (D) for fixed (F) and extracted neurons. C) Change in the average intensity values measured for secondary antibody signal along extracted neuron MTs, calculated by dividing the average intensity signal measured along extracted neuron axonal MTs or dendrites by that of fixed neuronal MTs. D) Ratio of change in the average intensity signal of acetylated tubulin along neuronal MTs relative to the change in tyrosinated tubulin signal, calculated by dividing the change in average intensity signal along extracted versus fixed axonal and dendritic MTs by that of the tyrosinated intensity signal. Data from 19-22 processes and 19-22 neurons from N = 3 independent experiment.

**Figure S8.**
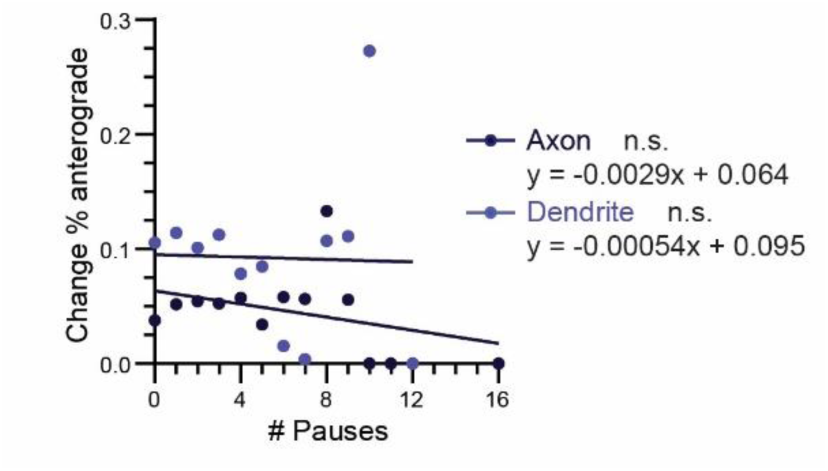
Motor pausing along extracted dendritic MT arrays is not correlated with track switches. Correlation between number of pauses and change in percentage of retrograde segments. Data was fitted with simple linear regression lines. F test (n.s. p > 0.05). Data from 59-80 processes from n = 49-50 neurons and N = 4 independent experiments.

**Figure S9.**
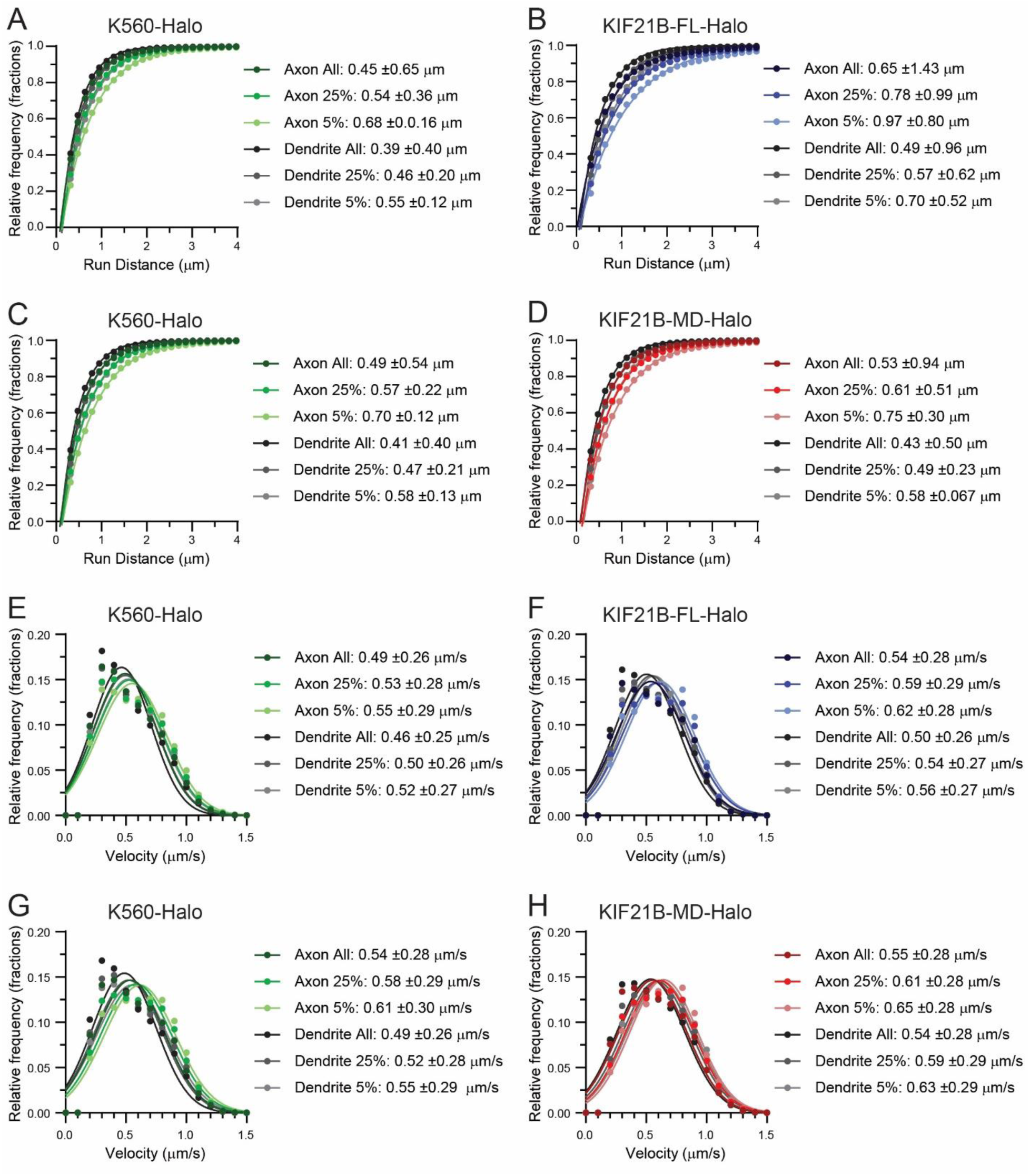
Kinesin motility is affected by MT arrangement and motor multimerization. A-D) Cumulative distribution of total distance K560-GFP and KIF21B-FL-Halo or KIF21B-MD-Halo motors traveled along extracted MT arrays in axons and dendrites. Data points were fitted by a single exponential decay function. Listed are the average distance traveled, as calculated by taking the inverse of the rate constant, and standard deviations. E-H) Histogram of velocities of motors along different MT bundles. Data points were fitted by a gaussian function. Listed are the means and standard deviations. Data from 24-42 processes from n = 21-28 neurons and N = 4 independent experiments.

**Figure S10.**
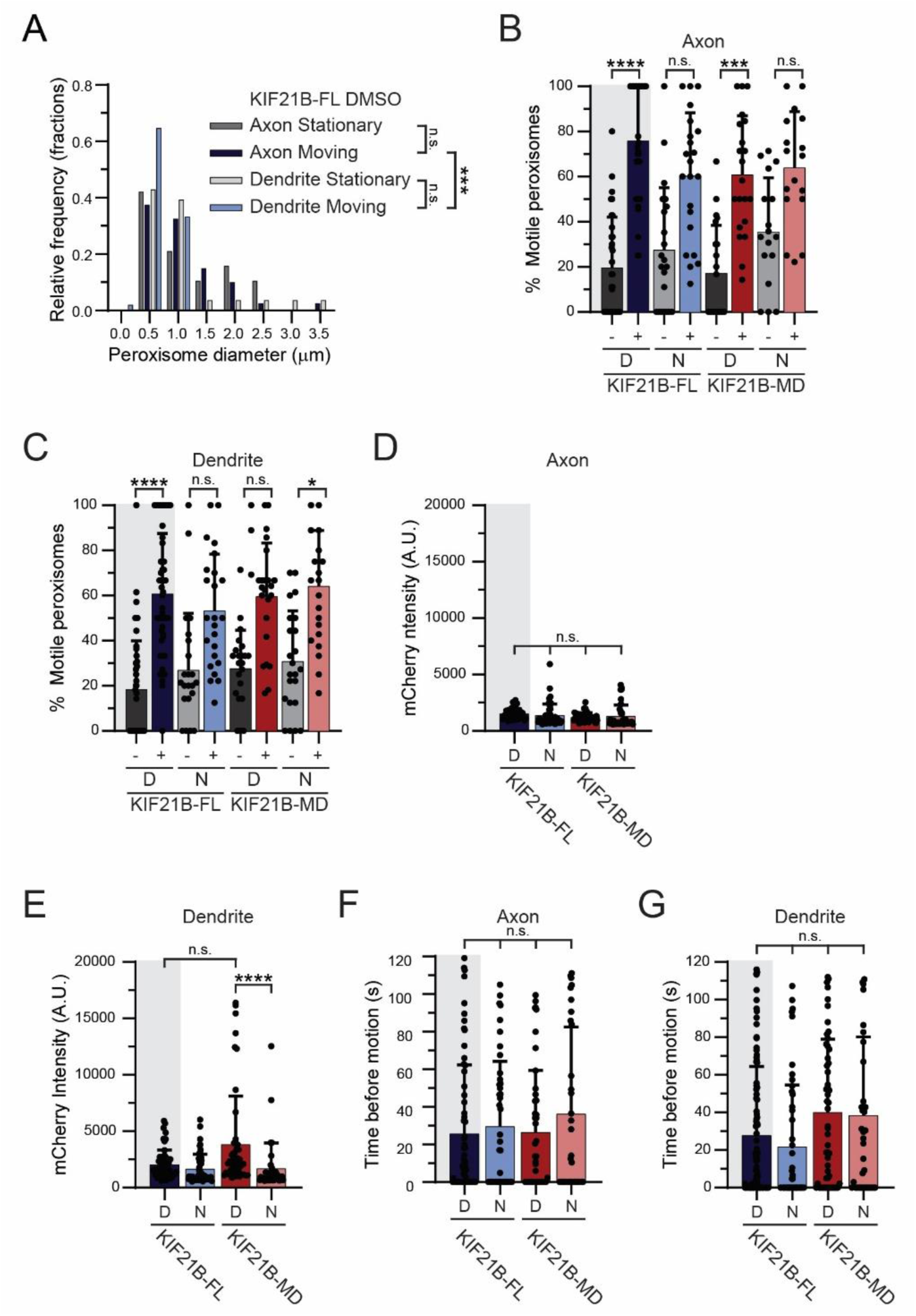
Induced recruitment of KIF21B motors to peroxisome cargo in live neurons promotes cargo movement. A) Histogram of KIF21B-FL recruited peroxisome diameter measured along axis of axons or dendrites that move post photoactivation or remain stationary in DMSO conditions. Kruskal–Wallis one-way ANOVA and Dunn’s multiple comparison (n.s. p > 0.05; ***p < 0.001). Data from 40-51 peroxisomes from n = 16-23 neurons and N = 3 independent experiments. B and C) Percentage of peroxisomes that are motile in axons and dendrites with and without photoactivation. Data for KIF21B-FL DMSO is repeated here from Figure 5 for comparison and marked with gray shading. DMSO (D) or nocodazole (N) conditions are indicated at the bottom of the graph. Plotted are means and standard deviations. Kruskal–Wallis one-way ANOVA and Dunn’s multiple comparison (n.s. p > 0.05; *p < 0.05; ***p < 0.001; ****p < 0.0001). D and E) Intensity of mCherry signal on moving peroxisomes with recruitment of KIF21B-FL or -MD. DMSO (D) or nocodazole (N) conditions are indicated at the bottom of the graph. Means and standard deviations are plotted. Kruskal–Wallis one-way ANOVA and Dunn’s multiple comparison (n.s. p > 0.05; ****p < 0.0001). F and G) Time taken by peroxisomes to begin movement post photoactivation in axons and dendrites. Data for KIF21B-FL DMSO is repeated here from Figure 5 for comparison and marked with gray shading. DMSO (D) or nocodazole (N) conditions are indicated at the bottom of the graph. Means and standard deviations are reported. Kruskal–Wallis one-way ANOVA and Dunn’s multiple comparison (n.s. p > 0.05). Unless otherwise stated, data from 37-120 peroxisomes from n = 18-52 neurons and N = 3-6 independent experiments.

**Figure S11.**
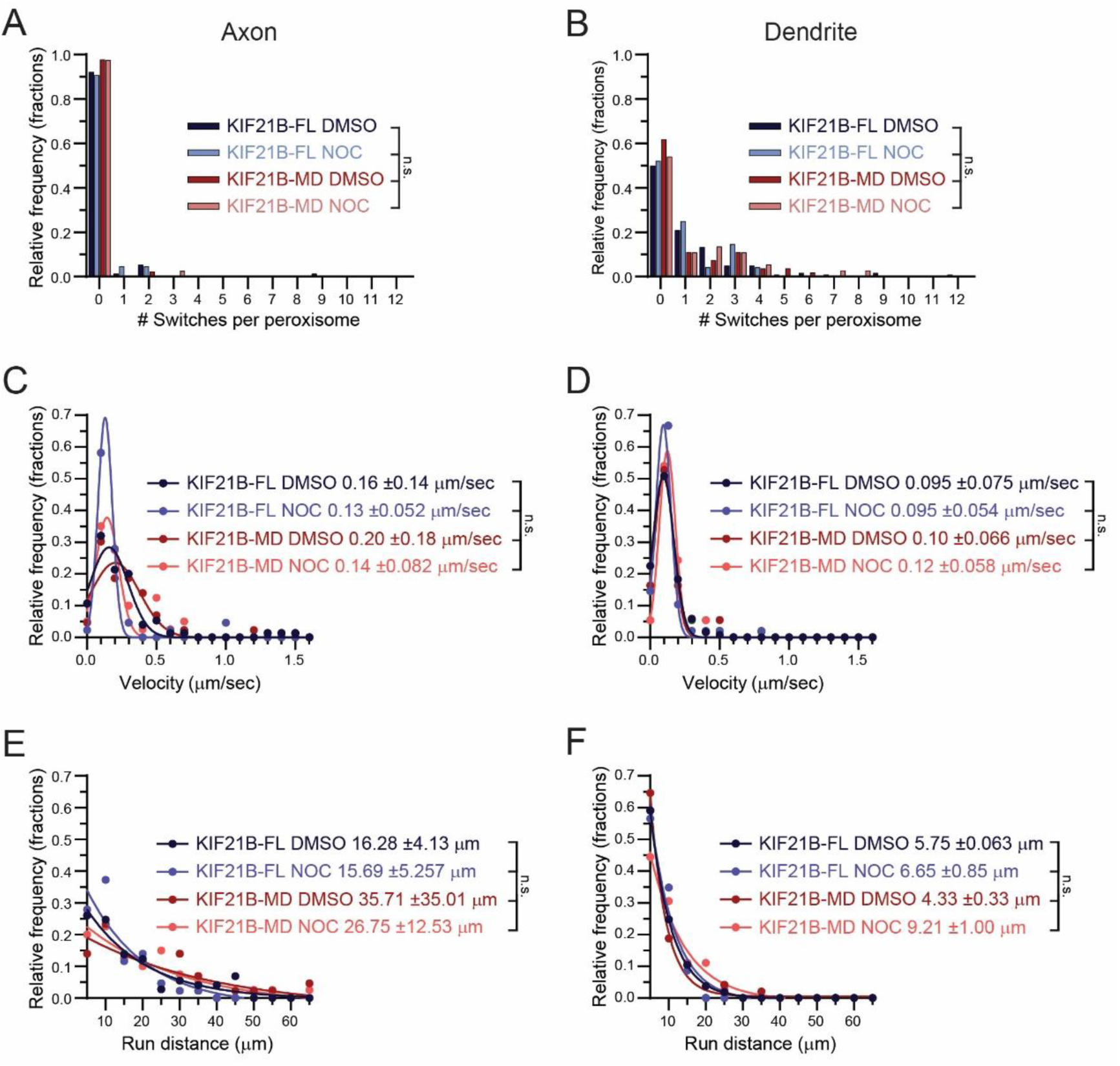
KIF21B induced retrograde trafficking in dendrites requires C-terminal tail domains and MT dynamics. A and B) Histogram of the number of track switches per peroxisome run in axons and dendrites. Motor and DMSO or nocodazole (Noc) conditions are indicated in the graph legend. Track switches are characterized by a reversal in direction that exceeds 0.4 µm in both anterograde and retrograde directions. Kruskal–Wallis one-way ANOVA and Dunn’s multiple comparison (n.s. p > 0.05). C and D) Histogram of velocities of motile peroxisomes after photoactivation in axons and dendrites. Motor and DMSO or nocodazole (Noc) conditions are indicated in the legend. Data points were fitted by a gaussian function. Listed are the means and standard deviations for each condition. Kruskal–Wallis one-way ANOVA and Dunn’s multiple comparison (n.s. p > 0.05). E and F) Histogram of run distances of motile peroxisomes after photoactivation. Motor and DMSO or nocodazole (Noc) conditions are indicated in the legend. Data points were fitted by a single exponential decay function. Listed for each condition is the average distance traveled, as calculated by taking the inverse of the rate constant, and standard deviations. Kruskal–Wallis one-way ANOVA and Dunn’s multiple comparison (n.s. p > 0.05). Data for KIF21B-FL DMSO is repeated here from Figure 5 for comparison. Unless otherwise stated, data from 37-120 peroxisomes from n = 18-52 neurons and N = 3-6 independent experiments.

## VIDEO LEGENDS

**Video 1. Purified KIF21B-FL motor motility along single dynamic MTs.** KIF21B-FL-Halo motors move towards the growing plus-end of the MT. Movie formatted at 30 FPS. Scale bar: 5 µm.

**Video 2. Purified KIF21B-FL motor motility along dynamic MTs forming a parallel bundle.** KIF21B- FL-Halo motors move towards the growing plus-ends of each MT. Movie formatted at 30 FPS. Scale bar: 5 µm.

**Video 3. Purified KIF21B-FL motor motility along dynamic MTs forming an antiparallel bundle.** KIF21B-FL-Halo motors move towards the growing plus-ends of each MT. Some motors switch from movement along one MT to the other to change direction. Movie formatted at 30 FPS. Scale bar: 5 µm.

**Video 4. Purified K560-GFP motor motility along extracted axonal MT arrays.** K560-GFP motors move mainly in the anterograde direction along axonal MTs. Movie formatted at 10 FPS. Scale bar: 5 µm.

**Video 5. Purified K560-GFP motor motility along extracted dendritic MT arrays.** K560-GFP motors move in both anterograde and retrograde directions along dendritic MTs. Some areas contain motor movement in mainly one direction. Movie formatted at 10 FPS. Scale bar: 5 µm.

**Video 6. Induced recruitment of KIF21B motors to peroxisome cargo in axons of live neurons.** KIF21B-FL recruited peroxisomes move in the anterograde direction immediately after photoactivation. White box indicates area of photoactivation. Green and red arrowheads indicate the locations of photoactivated peroxisomes with successful KIF21B-Fl recruitment in both the PEX3 and KIF21B fluorescence channel, respectively. Movie formatted at 10 FPS. Scale bar: 5 µm.

**Video 7. Induced recruitment of KIF21B motors to peroxisome cargo in dendrites of live neurons.** About 6 s after photoactivation, KIF21B-FL recruited peroxisomes in dendrites move in the retrograde direction. White box indicates area of photoactivation. Green and red arrowheads indicate the locations of photoactivated peroxisomes with successful KIF21B-FL recruitment in both the PEX3 and KIF21B fluorescence channel, respectively. Movie formatted at 10 FPS. Scale bar: 5 µm.

